# Development and validation of a multilocus sequence typing scheme for *Fasciola hepatica* using next-generation deep amplicon sequencing

**DOI:** 10.64898/2026.05.20.726500

**Authors:** Muhammad Abbas, Kezia Kozel, Olukayode Daramola, Nick Selemetas, Mark W. Robinson, Eric R. Morgan, Umer Chaudhry, Martha Betson

**Affiliations:** Discipline of Comparative Biomedical Sciences, School of Veterinary Medicine, University of Surrey, Guildford, UK; School of Veterinary Medicine, University of Lancashire, UK; Discipline of Microbes, Infection and Immunity, School of Veterinary Biosciences, University of Surrey, Guildford, UK; School of Biological Sciences, Queen’s University, Belfast, UK; Department of Veterinary Biomedical Sciences, Lewyt College of Veterinary Medicine, Long Island University, USA

**Keywords:** *Fasciola hepatica*, MLST, population genetics, SNP, polymorphism, Deep amplicon sequencing

## Abstract

Fasciolosis caused by *Fasciola hepatica* is an economically important disease in sheep and cattle. Knowledge of the population genetic structure of *F. hepatica* is important for understanding gene flow and informing disease control. In the present study, we designed, developed, and validated a multilocus sequence typing (MLST) scheme based on six markers. These markers were selected by aligning newly sequenced whole-genome sequence (WGS) data with available reference genomes and selecting variable regions with five or more single-nucleotide polymorphisms SNPs from different scaffolds of the *F. hepatica* reference genome *Fasciola* 10x pilon (GCA_900302435.1). Twenty markers were initially identified, of which 12 were multiplexed for deep amplicon sequencing after validation on worm and faecal eggs DNA; six markers were ultimately retained for downstream population genetics analysis. These markers were used to investigate population genetic structure in 15 cattle- and 27 sheep-derived *F. hepatica* populations in UK. A total of 53 unique alleles from six MLST markers were identified from 30 faecal (cattle = 13, sheep = 17) and 12 adult worm (cattle = 2, sheep = 10) populations. Shared alleles were observed in sheep- and cattle-derived populations. The highest allelic variation was observed in the Scottish Borders, Southern Scotland, and South-West England, and the lowest in North-West England. Minimal genetic differentiation was observed between cattle- and sheep-derived populations, with most genetic structuring within rather than between populations. Five markers showed high allelic polymorphism, whereas one marker showed low levels of allelic polymorphism, highlighting the importance of multilocus approaches. Overall, this six MLST-marker panel provides a tool for population genetic studies, revealing high gene flow and clonal expansion of *F. hepatica* across hosts and regions in the UK.

## Introduction

The liver fluke *Fasciola* parasite causes the economically important disease fasciolosis. The parasite infects a range of grazing animals, including cattle and sheep, as well as humans (Mas-Coma et al., 2018) and horses (Sanz et al., 2025). It poses a significant challenge to animal health and productivity (Charlier et al., 2014; Utrera-Quintana et al., 2022), and has a complex lifecycle (Calvani and Šlapeta, 2021; Vyas et al., 2026).

The intermediate hosts in this lifecycle are aquatic snails: *Galba truncatula* (Novobilský et al., 2014). Eggs are released in faeces, become embryonated, and hatch into miracidia that infect snails. Miracidia undergo asexual clonal expansion within the snail intermediate hosts, which then release cercariae into the environment (Dreyfuss and Rondelaud, 1994). One clonally-infected snail can produce up to 3000 clonal cercariae during its lifespan (Hodgkinson et al., 2018), contributing to pasture contamination and to the subsequent infection of hosts by metacercariae. This clonal expansion may also lead to a genetic bottleneck effect, particularly when infection levels in snail populations are low (Beesley et al., 2017). There is a strong possibility of the spread and random mixing of the metacercarial stage of parasites during the aquatic stage before infection of the definitive host, which can promote random mating in the final host (Criscione and Blouin, 2006). Small and large ruminants become infected by grazing on pasture contaminated with metacercariae (Dalton, 2021). A recent report showed metacercariae can survive for 10 weeks in contaminated silage under aerobic conditions and for two weeks under anaerobic conditions (John et al., 2020). *F. hepatica* is a hermaphroditic parasite and capable of both self and cross-fertilisation in the definitive host (Beesley et al., 2017) and can also reproduce asexually (Fletcher et al., 2004). These characteristics can impact allele frequency in populations.

The study of population genetic structures is essential for understanding parasite-host adaptation and the spread of traits such as drug resistance and adaptation to the host environment (Cwiklinski et al., 2015a, 2015b). Most population genetics analyses in *F. hepatica* are focused on mitochondrial markers including mt-ND1 and mt-COX1 (Elliott et al., 2014; Rehman et al., 2020; Semyenova et al., 2006). Mitochondrial DNA is haploid (Ballard and Whitlock, 2004) and has highly conserved regions (Ladoukakis and Zouros, 2017), maternal inheritance (Schwartz and Vissing, 2002), and is easy to amplify due to having a high copy number (Bogenhagen and Clayton, 1974). Mitochondrial loci are suitable for overview population genetic analysis, but they do not always act as strictly neutral loci (Ballard and Whitlock, 2004; Galtier et al., 2009). Their capacity is limited in resolving fine-scale population genetic structures (Zink and Barrowclough, 2008). Therefore, relying solely on mitochondrial DNA has limitations.

Nuclear microsatellite markers have been developed for *F. hepatica*, initially a panel of six microsatellite markers of which five were polymorphic (Hurtrez-Boussès et al., 2004). Later a multiplexed PCR panel of 15 highly polymorphic microsatellite loci was developed based on genome-derived data (Cwiklinski et al., 2015a). Using these microsatellite markers studies reported random mating and clonal transmission of parasite (Vázquez et al., 2016; Beesley et al., 2017, 2021), high genetic diversity (Beesley et al., 2017; Hecker et al., 2025), parasite clustering in different provinces in Argentina (Beesley et al., 2021) and high gene flow among dairy cattle farms in Germany (Hecker et al., 2025).However, in microsatellite markers, there can be high mutations and elevated levels of polymorphism and homoplasy (Hodel et al., 2016). Further, the large number of alleles per locus in microsatellites can inflate F-statistic values (Whitlock, 2011). On the other hand, allele-frequency distributions based on microsatellite markers can sometimes mislead the population genetic diversity because of fixation index (F_ST_) when the most common allele occurs at either very low or very high frequencies (Jakobsson et al., 2013). In addition, microsatellite datasets are prone to genotyping errors, which can bias downstream population genetic analyses (Hoffman and Amos, 2005).

To overcome the limitations of microsatellite typing, a genome-wide multilocus sequence typing (MLST) scheme can be an alternative approach to study the population genetics in *F. hepatica* (Schwabl et al., 2020). MLST schemes have recently been developed for protozoa, including *Cryptosporidium parvum* (Troell et al., 2025) and *Trypanosoma cruzi* (Diosque et al., 2014; Schwabl et al., 2020). Next-generation sequencing approach can amplify MLST alleles containing SNPs with high coverage. In general, multiplexed deep amplicon sequencing enables the simultaneous amplification of tens to hundreds of loci, even from samples with low DNA quantities, and bypassing the need for culturing or cloning (Peng et al., 2015; Lerch et al., 2017; Schwabl et al., 2020; Šlapeta et al., 2025).

This study aimed to develop a genome-wide MLST scheme for *F. hepatica* based on nuclear markers and apply this scheme to study the parasite’s population genetic structure including allelic variations, and gene flow across different regions of the UK.

## Methods

### Parasite and Egg Samples

This study utilised 86 *F. hepatica* positive samples, including 74 faecal egg samples and 12 adult worm samples. Collection and storage of these samples have been described in detail elsewhere (Abbas et al., 2026). The samples are grouped into populations by host species including cattle (n = 15), and sheep (n = 27) summarised in Supplementary Table 1.

### DNA Extraction and whole-genome sequencing

DNA was extracted from adult *F. hepatica* flukes based on morphological characteristics, as well as from faecal sedimented eggs, following the procedures described (Abbas et al., 2026). The extracted DNA was quantified and checked for quality using a BioDrop reader (Biochrom, UK) and Qubit dsDNA HS and BR Assay Kits (Invitrogen, USA). All DNA samples were stored at –80 °C until further analysis.

Genomic DNA from a single *F. hepatica* worm isolated from a cattle liver from West Sussex (sample ID: VPC-1) and from a worm collected from a sheep liver in County Tyrone (sample ID: MR-1) was submitted to Novogene (UK) for whole-genome sequencing (WGS) with 30X coverage. Samples VPC-1 and MR-1 were selected for WGS based on their high-quality genomic DNA (OD_260/280_ ratios of 1.92 and 1.83) and quantities (135 ng/µl and 66 ng/µl) (Supplementary Table 5). Briefly, genomic DNA was randomly fragmented, end-repaired, A-tailed, and ligated with Illumina sequencing adapters. Adapter-ligated fragments were size-selected, PCR-amplified, and purified to construct sequencing libraries. Library quality and concentration were assessed using Qubit fluorometric quantification and real-time PCR. The quantified libraries were pooled and sequenced using paired-end 150 bp (PE150) chemistry on an Illumina platform.

### Genome mapping and primer design for MLST

Illumina short reads (PE150) were extracted for two samples using the “tar -xvf” command on the High-Performance Cluster system at the University of Surrey, UK. The rFastp program in R version 4.3.0 was used for standard quality control assessment. A QC run was conducted on paired-end fastq files, including adapter trimming. Subsequently, both paired-end reads were merged into a single fastq file using rFastp (Chen et al., 2018).

Reference genomes of *F. hepatica* were downloaded from the NCBI WGS database projects (CANUEZ04, OMOY01, JXXN02) in .fna format. WGS data was aligned to reference sequences using Bowtie2 v2.5.1. First, Bowtie 2 index files were generated from the .fna genome file using the “*bowtie2-build*” command. The merged fastq file was aligned to reference genomes from the UK (GenBank assembly GCA_900302435.1, GCA_948099385.1 (replaced, GCA_948099385.2)) and the USA (GenBank assembly GCA_002763495.2) using Bowtie2 (Langmead et al., 2019; Langmead and Salzberg, 2012). The Bowtie 2 index command “*genome_index*” was used for alignment. The resulting alignment output files were saved in the “*output.sam*” file, and the alignment process utilised four threads. Overall alignment rates and mapping percentages were evaluated using Rsamtools in R version 4.3.0 (Morgan et al., 2016).

From the SAM files, binary files (BAM files) were generated using SAMtools-1.7 in bash through the “*samtools view -bS*” command. Subsequently, a pileup file was generated from the .bam file using the “*samtools mpileup*” command, followed by sorting with “*samtools sort*” and indexing the .bam file with “*samtools index*” in SAMtools. A Variant Call Format (VCF) file was generated using BCFtools-1.17 and the “*bcftools call -c -v -o*” command from both samples, followed by filtering and passing variants with GATK 4.4.0.0 (Bathke and Lühken, 2021; McKenna et al., 2010). Passed VCF files from all samples were merged using the command “*bcftools merge*” and a consensus sequence was generated using the “*bcftools consensus -f*” command. A BED file was produced in BCFtools using a threshold variable of two to six or more consecutive SNPs based on their positions in the merged VCF file. A BED file was used to extract 200 bp fragments (100 bp flanking regions from each side of identified SNPs) with the command “*bedtools getfasta -fi*” from different scaffolds using BEDTools-2.31.0. All R code scripts and bash command lines developed in this work can be accessed from the GitHub repository https://github.com/drmuhammadabbas810/New-tools-for-sustainable-control-of-liver-fluke-in-ruminants.git and Mendeley Data (https://data.mendeley.com/datasets/gc8fwkxdhy/1).

The FASTA files for the extracted fragments containing consecutive or non-consecutive five or more SNPs were visualised in Geneious version 8.0.5 (https://www.geneious.com) and verified using the UK scaffold level reference sequence (GCA_900302435.1) and 20 primer pairs (Supplementary Table 2) were designed to amplify each fragment using *Primer 3* parameters (Product size = 175 to 204, Primer size = 18 to 27 optimal= 20, Tm = 52 to 63 optimal= 60, GC%= 20 to 70% optimal= 50, Max temp difference = 2, Max Dimer tm= 47, Max 3/ stability = 9). Max Dimer Tm value of 47°C is a sensible choice to prevent the formation of stable primer dimers that could lead to non-specific amplification. It is within the typical range for dimer Tm values and helps maintain primer specificity. A Max 3 /Stability value of 9 indicates that the 3’ ends of the primers should not form highly stable secondary structures with each other. Figure 1 summarises the bioinformatics workflow for marker identification.

**Figure 1.**
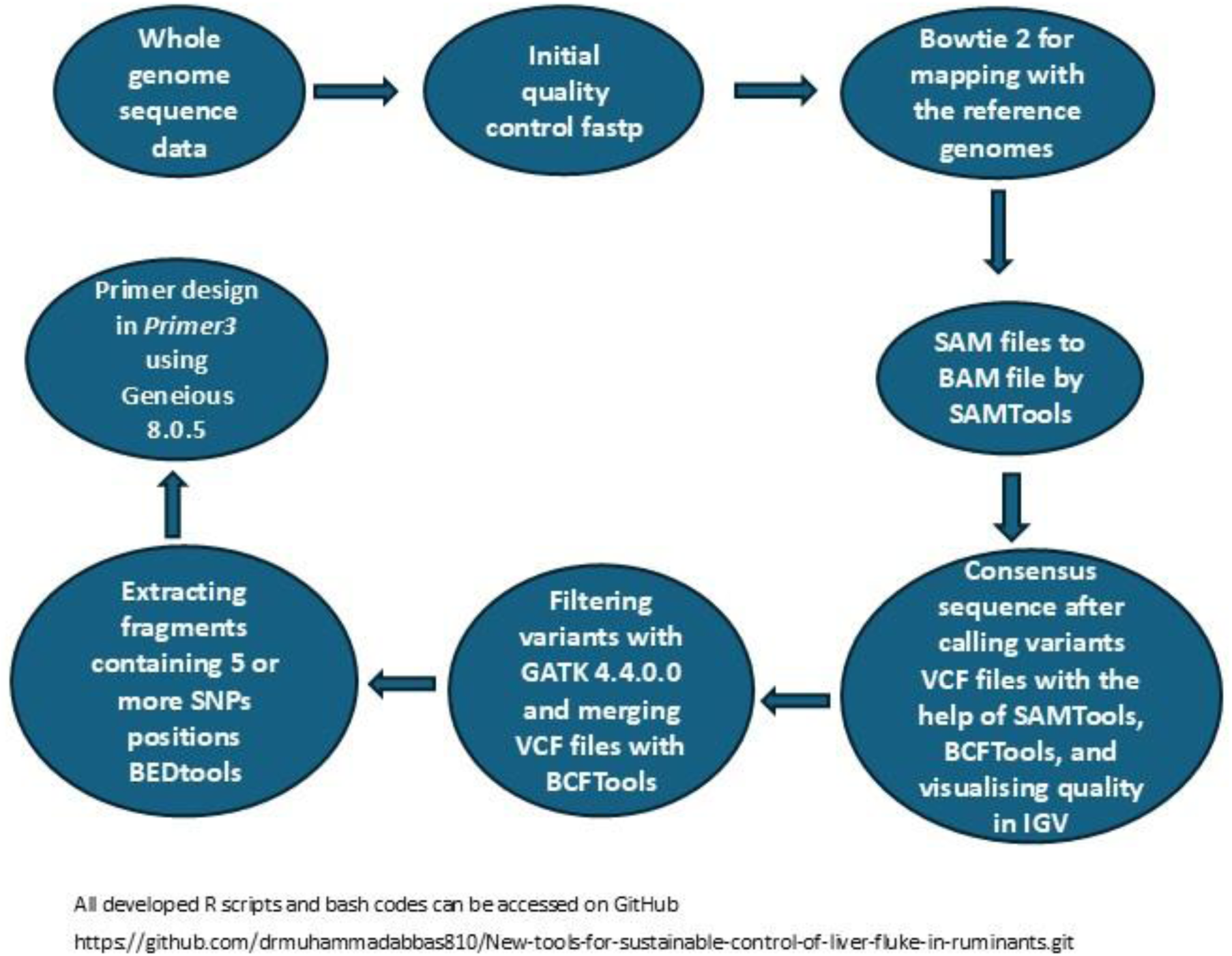
Bioinformatics workflow for the development of MLST markers

### MLST marker selection and testing

Two to six SNP-containing fragments ranging from four to more than a hundred thousand were extracted from three *F. hepatica* reference genomes for the choice of polymorphic regions (Table 1). Using the criterion of consecutive or non-consecutive SNPs, the UK reference genome GCA_900302435.1 (Fasciola_10x_pilon) showed informative fragments, including five or more SNPs (n = 25) and four or more SNPs (n = 207). The second UK genome, GCA_948099385.1 (FhHiC23), showed slightly fewer SNP-rich regions, producing 22 fragments with ≥5 SNPs and 181 fragments with ≥4 SNPs. In comparison, the USA reference genome GCA_002763495.2 (F_hepatica_1.0.allpaths) produced 24 fragments containing ≥5 SNPs, and the highest number of fragments containing ≥4 SNPs (n = 243).

**Table 1.** Summary of consecutive and non-consecutive SNP rich fragments identified in available reference genomes used for MLST marker selection.

| Reference genome | Fragments with 6 or more SNPs | Fragments with 5 or more SNPs | Fragments with 4 or more SNPs | Fragments with 3 or more SNPs | Fragments with 2 or more SNPs |
| --- | --- | --- | --- | --- | --- |
| GCA_900302435.1Fasciola_10x_pilon (UK)* | 5 | 25 | 207 | 3307 | 99,769 |
| GCA_948099385.1_FhHiC23_genomic.fna (UK) | 4 | 22 | 181 | 3051 | 94,459 |
| GCA_002763495.2_F_hepatica_1.0.allpaths (USA) | 4 | 24 | 243 | 4340 | 1,28,518 |

\*Primers were designed based on extracted fragments containing 5 or more SNPs from the reference genome GCA\_900302435.1Fasciola\_10x\_pilon (UK)

When comparing lower SNP thresholds, all genomes produced substantially higher fragment counts. However, based on comparative SNP presence and fragment consistency, and a visual inspection of fragments, 20 primers were initially designed from the set of unique fragments containing ≥5 SNPs extracted from the UK reference genome GCA_900302435.1 (Fasciola_10x_pilon).

All 20 designed MLST primer pairs (Supplementary Table 2) were tested to determine whether they produced amplicons using positive-control genomic DNA from adult worms and faecal eggs. PCRs were performed in 25 µL reactions containing 12.5 µL DreamTaq Green PCR Master Mix (Thermo Scientific), forward and reverse primers (0.20 µM), 4 µL of genomic DNA and nuclease-free water. Thermocycling conditions consisted of an initial denaturation at 95 °C for 5 minutes, followed by 35 cycles of 95 °C for 1 minute, 59.9 °C for 1 minute, and 72 °C for 1 minute, with a final extension at 72 °C for 5 minutes. 2 µL of PCR product was visualised on a 2% agarose gel; the tested molecular markers generated bands ranging from 175 to 200 bp. Of these 20 primers, two primers failed to generate amplicons in PCR reactions and were excluded from further analysis. The remaining PCR mix containing amplicon products was purified using magnetic bead purification (MACHEREY-NAGEL GmbH & Co. KG). Purified products were Sanger-sequenced by Source Bioscience (UK). Resulting chromatograms were visualised and trimmed in Geneious v8.0.5. FASTA sequences of MLSTs were aligned to reference genome scaffolds to confirm locus identity.

The 18 MLST markers verified by sanger sequencing were metabarcoded and tested on DNA extracted from adult worms and randomly selected DNA samples extracted from *F. hepatica* eggs (EPG range: 4-121 SE: 23.99) from five UK counties (Cumbria, Scottish Borders, Peeblesshire, South Lanarkshire, and Dorset) that had previously tested positive for *F. hepatica* by deep sequencing and/or qPCR (Abbas et al., 2026). The 18 metabarcoded MLST markers showed PCR amplification success for each marker varied from 9 to 14 markers on genomic DNA isolated from different *F. hepatica* eggs obtained from five different regions of the UK and is summarised (Supplementary Table 7). Finally, the well performing 12 MLST markers were selected, including 3-327, 6-673, 8-1410, 9-1532, 10-1632, 11-1914, 12-2022, 14-2089, 15-2089, 17-2194, 19-2745, and multiplexed into two reactions for deep amplicon sequencing (Supplementary Table 3). PCR reactions were performed in 25 µL volumes consisting of 5 µL of KAPA HiFi fidelity buffer, 0.5 U KAPA HiFi fidelity polymerase (KAPA Biosystems, South Africa), 0.3 mM dNTPs, and pools of forward and reverse primers containing four forward and four reverse primers with a final concentration of 300 nM per MLST primer, along with 3 µL of DNA template and nuclease-free water. PCR cycling conditions were an initial denaturation at 95 °C for 2 minutes, followed by 35 cycles of 98 °C for 20 seconds, annealing at 62 °C for 15 seconds, and extension at 72 °C for 15 seconds, with a final extension at 72 °C for 5 minutes. Resulting PCR products containing amplicons were purified using magnetic bead purification (MACHEREY-NAGEL GmbH & Co. KG) and subjected to a second round of PCR for library preparation. The primer sets, adaptors, barcoded PCR amplification conditions, final magnetic bead purification, and final library quantification were conducted as described previously (Sargison et al., 2019; Yasein et al., 2022).

### Demultiplexing of MLST sequence reads and bioinformatics analysis

The Illumina MiSeq demultiplexed the raw sequencing data and generated corresponding FASTQ files for each sample by identifying the sample-specific barcoded indices after the run was complete (Supplementary Table 4; NCBI Bioproject: PRJNA1431397; Accession numbers SAMN56308754 to SAMN56308884 and SAMN56308979 to SAMN56309022; https://data.mendeley.com/datasets/gc8fwkxdhy/1). The FASTQ files were further demultiplexed for each MLST marker using Mothur versions 1.41.0 and 1.48.1 (Schloss et al., 2009) on the High-Performance Computing (HPC) cluster at the University of Surrey, UK. This processing followed pipelines described in previous studies (Rehman et al., 2020; Sargison et al., 2019) with modifications and pipelines available in the Mendeley Data repository (https://data.mendeley.com/datasets/gc8fwkxdhy/1).

For demultiplexing of each MLST locus, corresponding sequences were extracted using a reference library generated from the sequence fragments extracted from reference genomes GCA_900302435.1, GCA_948099385.1, GCA_002763495.2 and Sanger sequencing data (https://data.mendeley.com/datasets/gc8fwkxdhy/1). The Mothur pipeline joined paired-end reads, filtered out ambiguous or low-quality sequences, and removed reads that were excessively long or short. Unique and identical sequences were pre-clustered, and abundant reads were grouped to reduce sequencing errors and improve accuracy. Abundant alleles for each MLST locus were extracted from the filtered dataset using a sequence-read threshold of 1,000 to 10,000 reads per MLST marker to remove noise and sequencing artefacts. These thresholds were determined after visual inspection of the output count table files from Mothur analysis. Downstream processing was performed in R. Raw sequence reads from each sample were assigned to populations (Supplementary Table 1) and exported as FASTA files. To ensure sequence quality, all reads were cleaned by removing ambiguous nucleotides and retaining only sequences ≥100 bp. Cleaned sequence reads were saved as population-specific FASTA files for each MLST marker. For each population, sequences were replicated according to their read counts to generate read-weighted FASTA files. In parallel, unique sequences were extracted from FASTA files for each population and each MLST marker for downstream analyses.

### Data Analysis

All analyses were performed in R version 4.3.3 (https://cran.r-project.org/). Alleles were sorted by total abundance, and allele IDs were assigned in descending order. The threshold criteria for final marker selection were set at working on at least 60% of faecal egg populations, more than 90% of worm populations, and 10,000 sequence reads from all populations. To assess allelic variation, cattle and sheep allele sequences for each MLST marker were aligned using MUSCLE (Bodenhofer et al., 2015). SNPs were detected by identifying alignment sites with different nucleotides. Within each population of cattle and sheep, all alleles were aligned separately, and SNPs were counted for each allele in the populations, producing per-allele SNP counts and overall SNP statistics. Finally, all allele sequences for each marker were visualised using Geneious v8.0.0. All R scripts and the reference sequences library used in this work are available at the Mendeley data repository (https://data.mendeley.com/datasets/gc8fwkxdhy/1).

MLST data sets were assigned to populations based on host (sheep and cattle). Total sequence reads were summarised for sheep- and cattle-derived *F. hepatica* populations and further compared by sample source (worms vs. faeces). The proportion of sequence reads in different populations showing each MLST loci was calculated, and stacked percentage bar charts were produced to visualise MLST distributions across all populations using readxl (Wickham et al., 2025a) and tidyverse (Wickham et al., 2019).

Allele presence metrics were calculated at the population level for each host, including the number of alleles, mean alleles per population, and standard deviation. For each MLST marker, the number of alleles, total SNP sites, total SNPs, mean, and ± S.D. SNPs per allele were calculated separately for sheep and cattle using the following R packages: phylotools (Revell, 2024), microseq (Snipen and Liland, 2025), tidyverse (Wickham et al., 2019), dplyr (Wickham et al., 2023), tidyr (Wickham et al., 2025b), purrr (Wickham et al., 2026a), ggplot2 (Wickham, 2016), seqRFLP (https://cran.r-project.org/src/contrib/Archive/seqRFLP/), readxl (Wickham et al., 2025a), writexl (Ooms et al., 2025), ggh4x (Brand, 2025), Biostrings (Pagès et al., 2025), seqinr (Charif and Lobry, 2007), and msa (Bodenhofer et al., 2015).

Principal Coordinate Analysis (PCoA) was performed for the six selected MLST markers using the distance matrices from allele sequences of sheep- and cattle-derived populations. Pairwise distance matrices were generated and analysed using R packages readr (Wickham et al., 2026b), dplyr (Wickham et al., 2023), ggplot2 (Wickham, 2016), and cowplot (Wilke, 2025). Population clustering on PCoA plot was assessed by geographic region and host. AMOVA and pairwise F_ST_ values were calculated in Arlequin v3.5 (Excoffier and Lischer, 2010) to evaluate genetic differentiation within and between host-associated groups. Allelic sequence distribution across all populations was visualised using heatmaps generated with pheatmap (Kolde, 2025) and ggplot2 (Wickham, 2016), displaying both raw and log-transformed sequence read counts scaled by row to highlight relative sequence reads abundance patterns.

## Results

### Quality assessment of WGS data and mapping of reads to the reference genome

Illumina sequencing generated a total of 47 GB of sequencing short reads data. The raw sequencing data showed Q30 values of 93.99% for VPC-1 (accession no. SAMN56321788) and 94.57% for MR-1 (accession no. SAMN56321787), with GC contents of 44.30% and 44.24%, respectively. Following filtration, adapter trimming and merging of forward and reverse reads, samples VPC-1 and MR-1 showed Q30 base percentages of 81.0732% and 90.7411%, respectively, with duplication rates of 25.4014% and 24.8026%. For VPC-1, the overall alignment rate with the reference genomes ranged from 88.89% to 95.39%, and the mapping percentage ranged from 99.90% to 100%. For MR-1, the alignment rate ranged from 90.20% to 96.57%, and the mapping rate ranged from 99.91% to 100% (Supplementary Table 6).

### Performance of MLST markers

A total of 11 MLST markers generated sequencing reads (in total 4,682,733) from 12 worm populations (100%) (reads in total 1,044,132), and 72 egg samples (97.29%) across 30 different populations (reads in total 3,638,601). A total of 1,102,154 reads from eggs purified from cattle faecal populations (n= 13), 232,899 reads from cattle worm populations (n = 2), 2,536,447 reads from eggs purified from sheep faecal populations (n = 17), and 811,233 reads from sheep worm populations (n = 10).

Among the 11 MLST markers evaluated, sequence data were available for MLSTs 8-1410 and 14-2089 in 100% of worm samples and in over 83% of egg samples (see Table 2). Reads for MLSTs 3-327, 9-1532, 17-2194, and 19-2745 were detected for 91.67% to 100% of worm populations and 60% to 80% of egg populations. In contrast, MLSTs 6-673, 11-1914, 12-2022, 10-1632, and 20-2815 sequences were found in a smaller proportion of worm populations (83% for some markers) and a proportion of egg populations (<40%). The overall number of sequence reads was lower than 10,000 in total from all populations per marker for the latter markers. This suggested the limited suitability of these markers for population-wide genotyping, especially from egg populations. Based on these results, six MLST markers were finally selected for downstream population analyses: 3-327, 8-1410, 9-1532, 14-2089, 17-2194, and 19-2745. The presence of these markers in different populations is presented in Figure 2 and Supplementary Figure 1. A total of 4.5 million sequence reads were generated for 53 unique alleles from the six selected markers. After normalization, the sequence read counts for each allele in the 27 sheep- and 15 cattle-derived *F. hepatica* populations are summarised in a heatmap (Figure 3) and without normalisation in Supplementary Figure 2.

**Figure 2.**
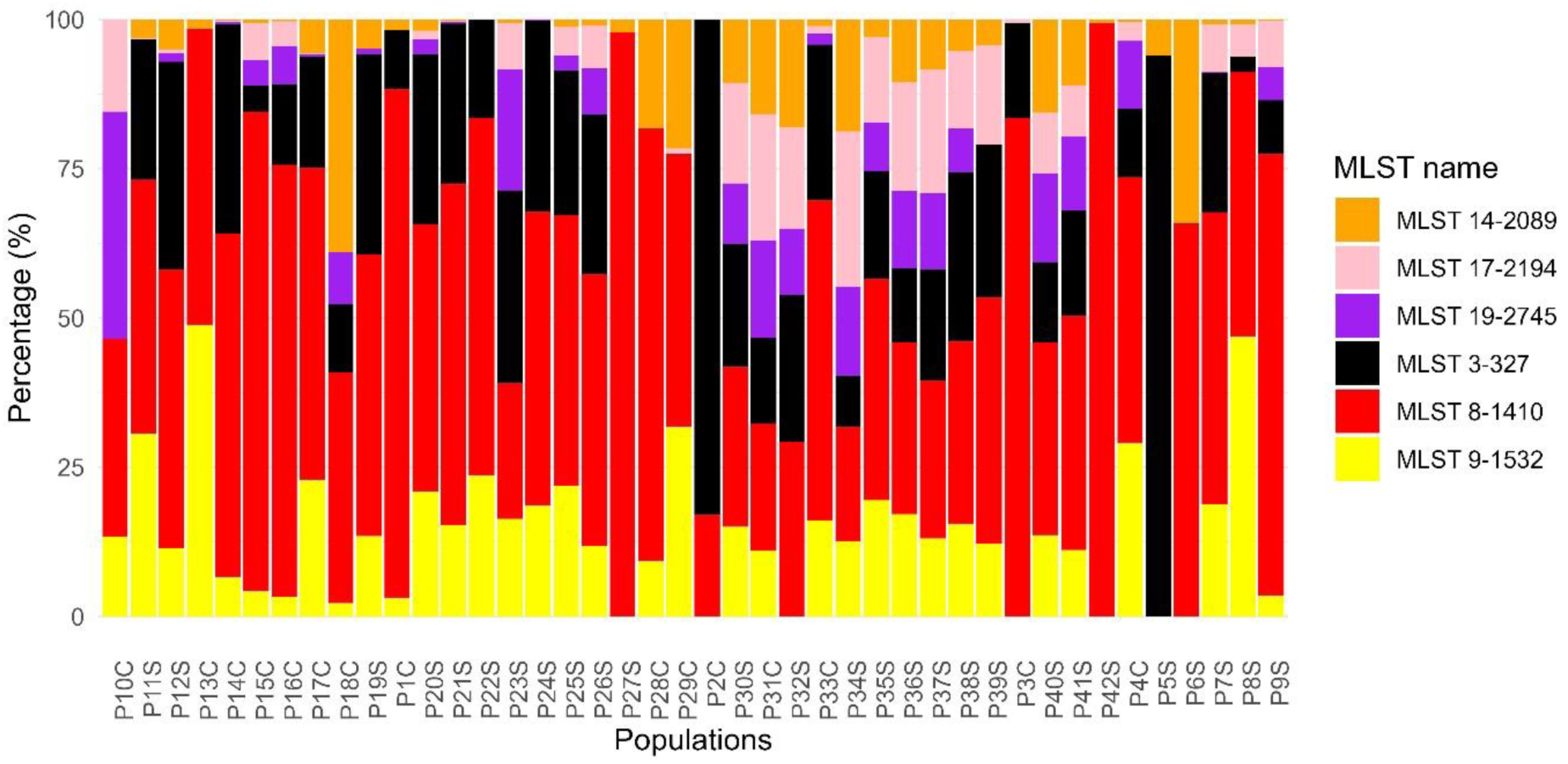
Distribution of sequence reads generated by six MLST markers across different populations. The X-axis represents populations from cattle (C) and sheep (S), while the Y-axis shows the percentage of sequence reads from each MLST marker.

**Figure 3.**
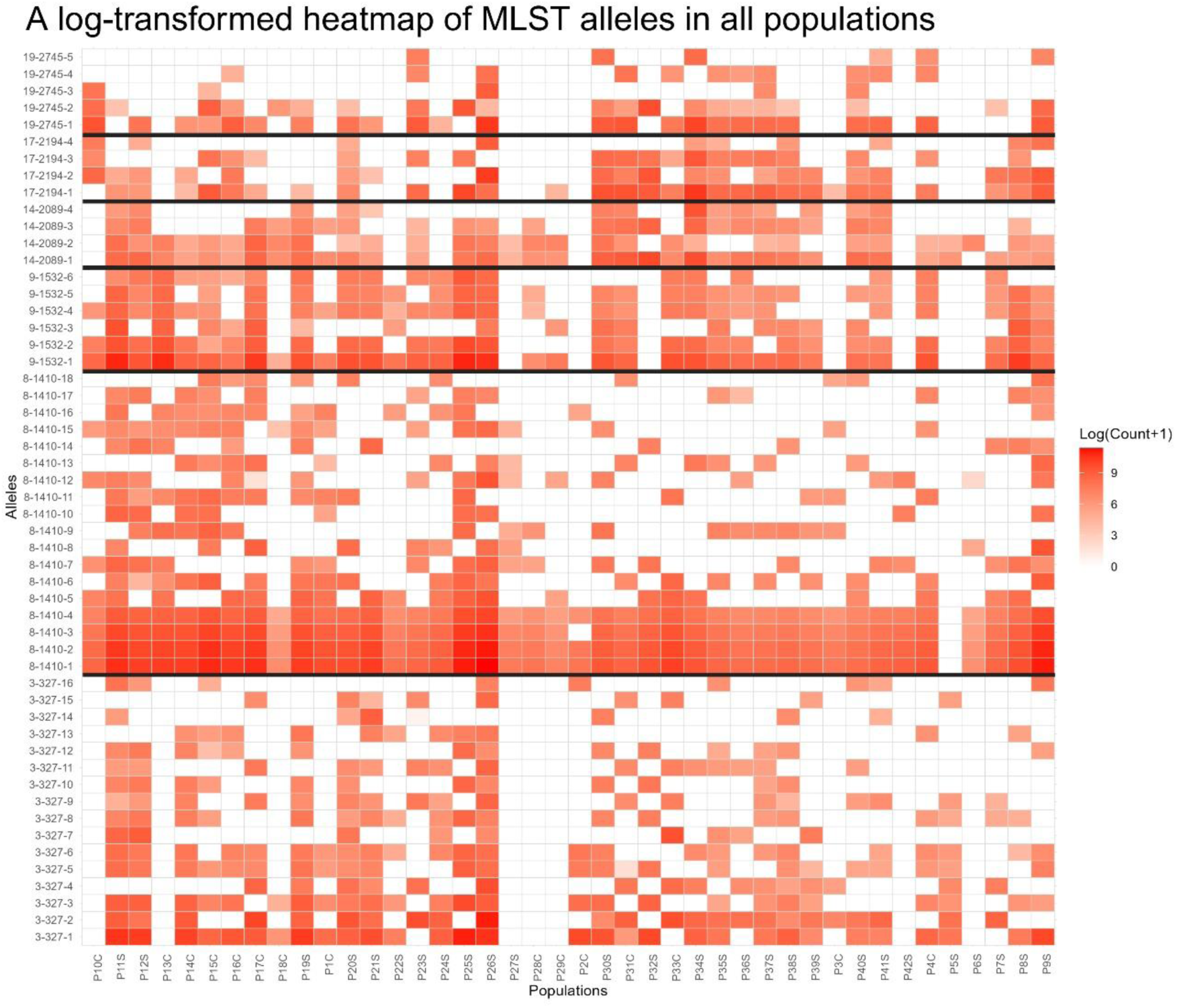
Heatmap displaying normalised sequence read counts for 53 alleles detected at six MLST marker loci in 27 sheep-derived and 15 cattle-derived *F. hepatica* populations. Colour intensity represents the relative abundance of each allele within each population. A black line separates alleles for each MLST.

**Table 2.** MLST marker performance in eggs and adult worms DNA of *F. hepatica*.

| MLST marker | No of egg populations generated sequence reads |  | No of worm populations generated sequence reads |  | Total cattle and sheep-derived populations generated sequence reads |  |  | Total populations tested |  | Cattle and sheep-derived populations generated reads in percentage (%) |  | Total sequence reads generated in all populations |
| --- | --- | --- | --- | --- | --- | --- | --- | --- | --- | --- | --- | --- |
|  | Cattle | Sheep | Cattle | Sheep | Eggs | Worms | Total | Eggs | Worms | Eggs | Worms |  |
| 3-327 | 9 | 14 | 2 | 10 | 23 | 12 | 35 | 30 | 12 | 76.67 | 100 | 952862 |
| 6-673* | 4 | 8 | 2 | 10 | 12 | 12 | 24 | 30 | 12 | 40 | 100 | 24682 |
| 8-1410 | 13 | 16 | 2 | 10 | 29 | 12 | 41 | 30 | 12 | 96.67 | 100 | 2182903 |
| 9-1532 | 11 | 13 | 2 | 9 | 24 | 11 | 35 | 30 | 12 | 80 | 91.67 | 742342 |
| 10-1632* | 0 | 1 | 0 | 5 | 1 | 5 | 6 | 30 | 12 | 3.33 | 41.67 | 5030 |
| 11-1914* | 1 | 2 | 2 | 8 | 3 | 10 | 13 | 30 | 12 | 10 | 83.33 | 23941 |
| 12-2022* | 1 | 2 | 1 | 4 | 3 | 5 | 8 | 30 | 12 | 10 | 41.67 | 46899 |
| 14-2089 | 10 | 15 | 2 | 10 | 25 | 12 | 37 | 30 | 12 | 83.33 | 100 | 178164 |
| 17-2194 | 8 | 11 | 2 | 10 | 19 | 12 | 31 | 30 | 12 | 63.33 | 100 | 280601 |
| 19-2745 | 7 | 11 | 2 | 9 | 18 | 11 | 29 | 30 | 12 | 60 | 91.67 | 238324 |
| 20-2815* | 0 | 1 | 0 | 3 | 1 | 3 | 4 | 30 | 12 | 3.33 | 25 | 6985 |

\*Markers excluded from downstream analysis based on a threshold either sequence reads 10,000 in total from all populations, or marker performance less than 60% using egg DNA. MLST 15-2089 did not produce any reads.

### Allelic variations in MLSTs

Substantial allelic variations were observed across MLST markers in different regions. Overall, sheep-derived populations showed higher allelic diversity than cattle-derived populations. Based on the number of alleles observed in the different regions of the UK, MLSTs 3-327 and 8-1410 were the most genetically diverse markers with high allele count in both host, whereas MLSTs 17-2194 and 19-2745 were the least allelic count loci. Populations from the Scottish Borders, Southern Scotland, and parts of South-West England showed the allelic richness, while populations from North-West England displayed the lowest allelic richness (Table 3).

**Table 3.**
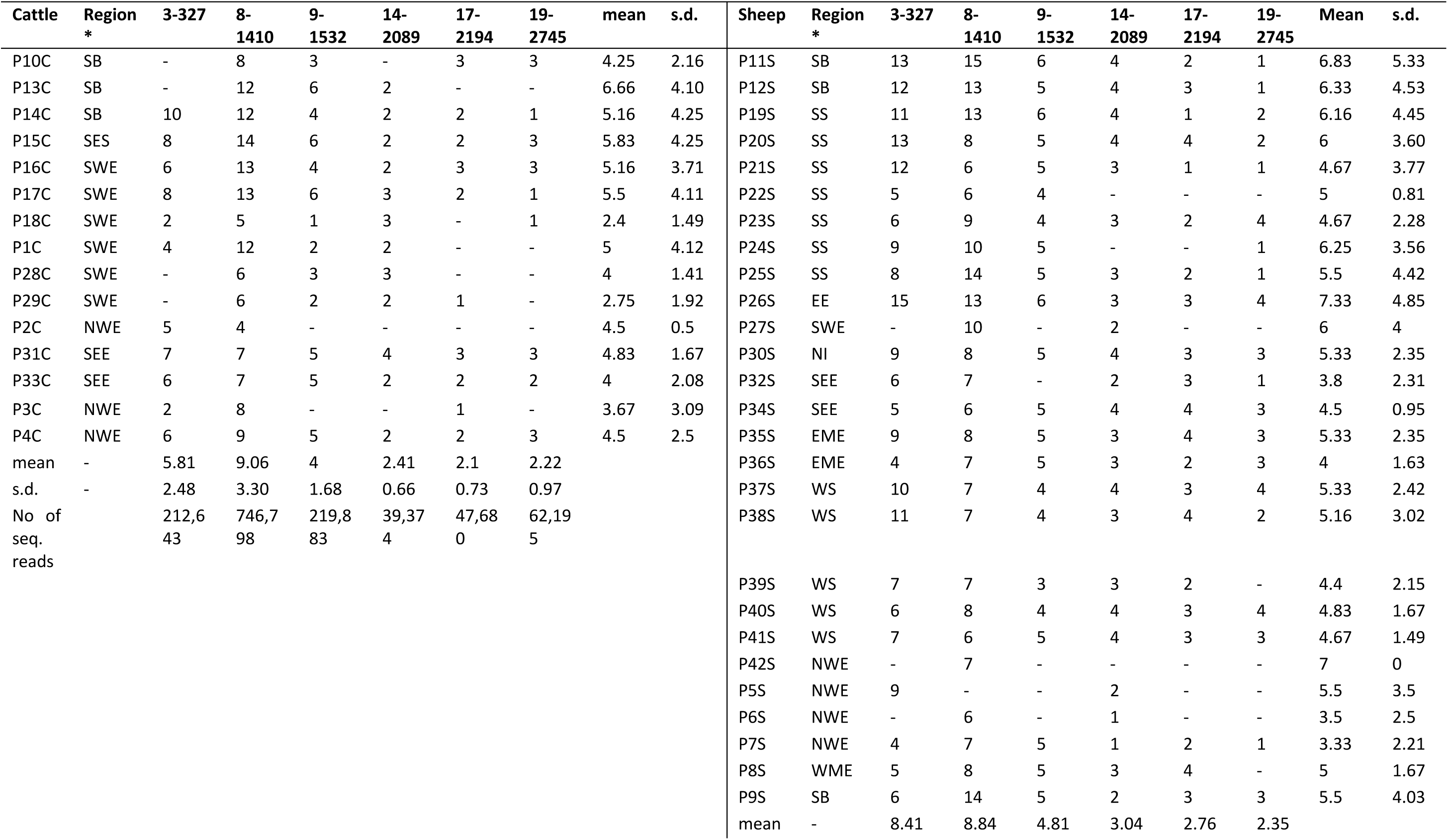
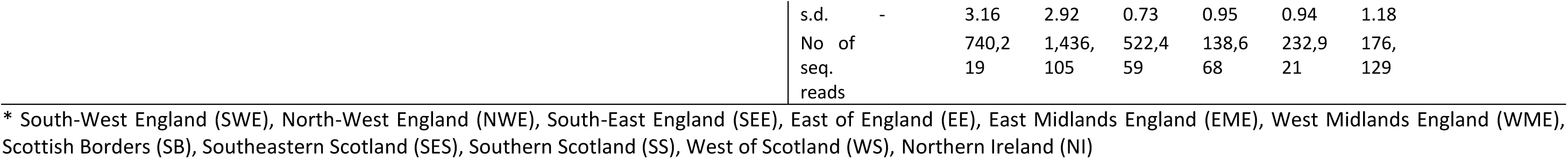
Number of alleles per population based on a panel of 6 MLST markers in 15 cattle and 27 sheep-derived *F. hepatica* populations. Each value in the data set shows the number of alleles found in each population.

In cattle-derived populations, the highest allelic counts were observed at MLST-8-1410, with a mean number (±SD) of 9.06 (±3.30), particularly in populations from the Scottish Borders and South-West England. MLSTs 3-327, 8-1410, 17-2194, and 19-2745 showed lower allele numbers, especially in populations from North-West England.

Sheep-derived populations showed high allele counts at MLSTs 3-327 and 8-1410 (mean = 8.41 ±3.16 and 8.84 ±2.92, respectively). Populations from the Scottish Borders and Southern Scotland exhibited high allelic means (mean = 4.67–6.83). Low allelic variations were observed in several North-West England populations (mean = 3.33 ± 2.21 and 7 ± 0.0), because multiple loci had no alleles. The highest allelic means were observed in a single population from the East of England (mean = 7.33 ± 4.85), which also showed the highest allelic counts across all loci (Table 3).

The relationship between sequencing read depth and the number of alleles detected in six MLST markers was analysed for *F. hepatica* populations from cattle and sheep. Spearman’s rank correlation coefficient (Spearman’s *ρ*) indicated a strong positive association for both host species. For cattle, *ρ = 0.93* (*P = 0.008*; *R² = 0.72*), and for sheep, *ρ = 0.90* (*P = 0.015*; *R² = 0.84*) were recorded, indicating that MLST loci with higher sequence reads result in a higher number of alleles (Figure 4).

**Figure 4.**
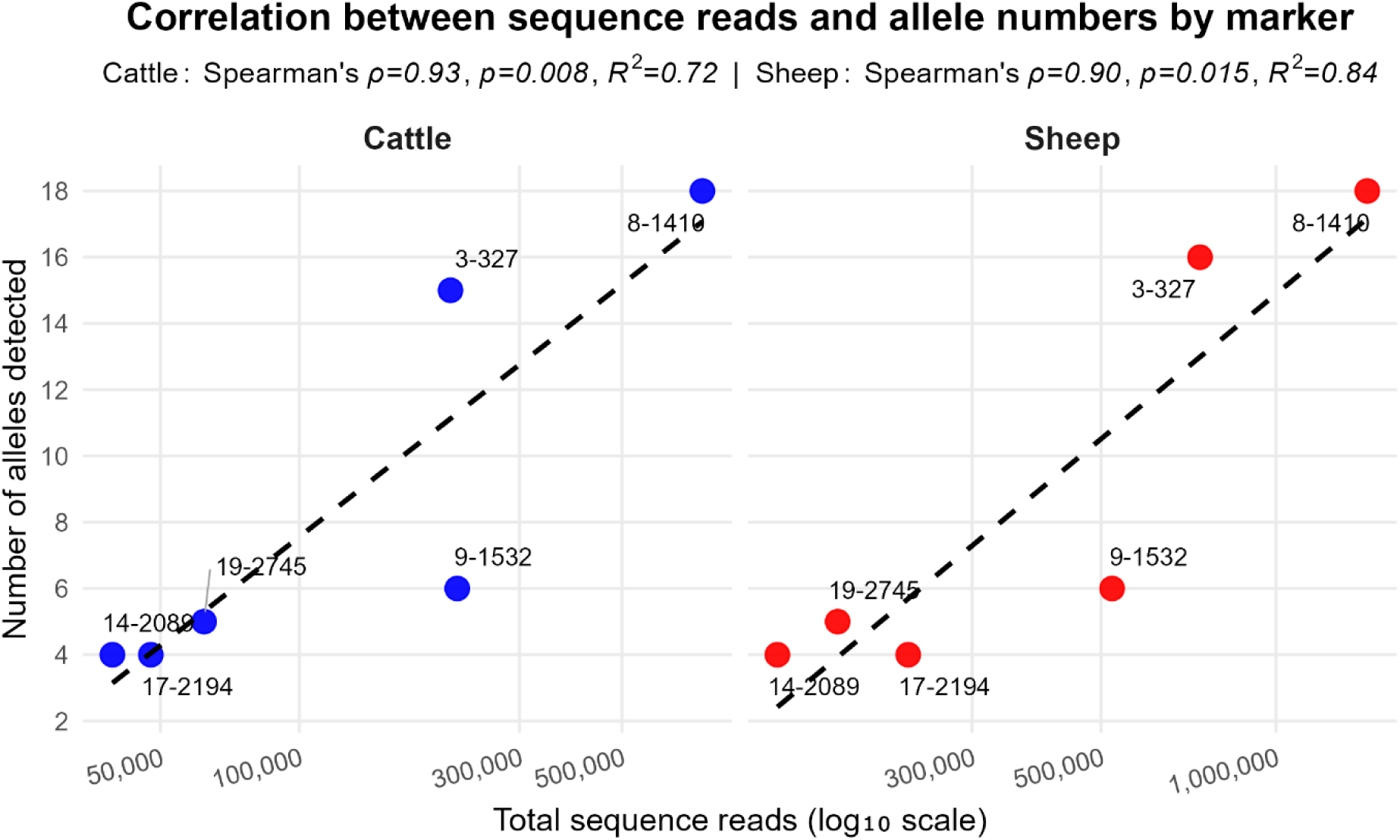
Correlation between the sequencing reads generation and number of alleles from six MLST markers in 15 cattle- and 27 sheep-derived *F. hepatica* populations in the UK. The Spearman’s rank correlation coefficient was recorded for cattle (ρ = 0.93, P = 0.008) and sheep (ρ = 0.90, P = 0.015) and R^2^ correlation was noted for cattle (R² = 0.72) and sheep (R² = 0.84). These values demonstrate that the allele numbers detected in different MLSTs are significantly correlated with sequencing read depth in cattle- and sheep-derived *F. hepatica* populations.

Allelic variation patterns are largely consistent between sheep and cattle host, suggesting the sharing of alleles between hosts (Figure 5, Table 4). The number of alleles per MLST locus ranged from 4 to 18. The highest alleles were observed at MLST 8-1410, followed by 3-327 in both cattle and sheep hosts. However, the 3-327 marker was more polymorphic, containing an average of 3.93 and 3.94 SNPs per allele in cattle and sheep-derived parasite populations, respectively, compared to 8-1410, which included an average of 2.06 SNPs in both host-derived *F. hepatica* populations. MLST 19-2745 and 17-2194 were also polymorphic, with five and four alleles, respectively, and an average of 2.4 and 2.5 SNPs per allele in cattle- and sheep-derived parasite populations. MLSTs 14-2089 showed moderate polymorphism (average 1.5 SNPs per allele), with 4 alleles in both host species. The MLST 9-1532 locus was monomorphic and highly conserved, with an average of only 0.17 SNPs per allele and six alleles were recorded in both hosts (Table 4).

**Figure 5.**
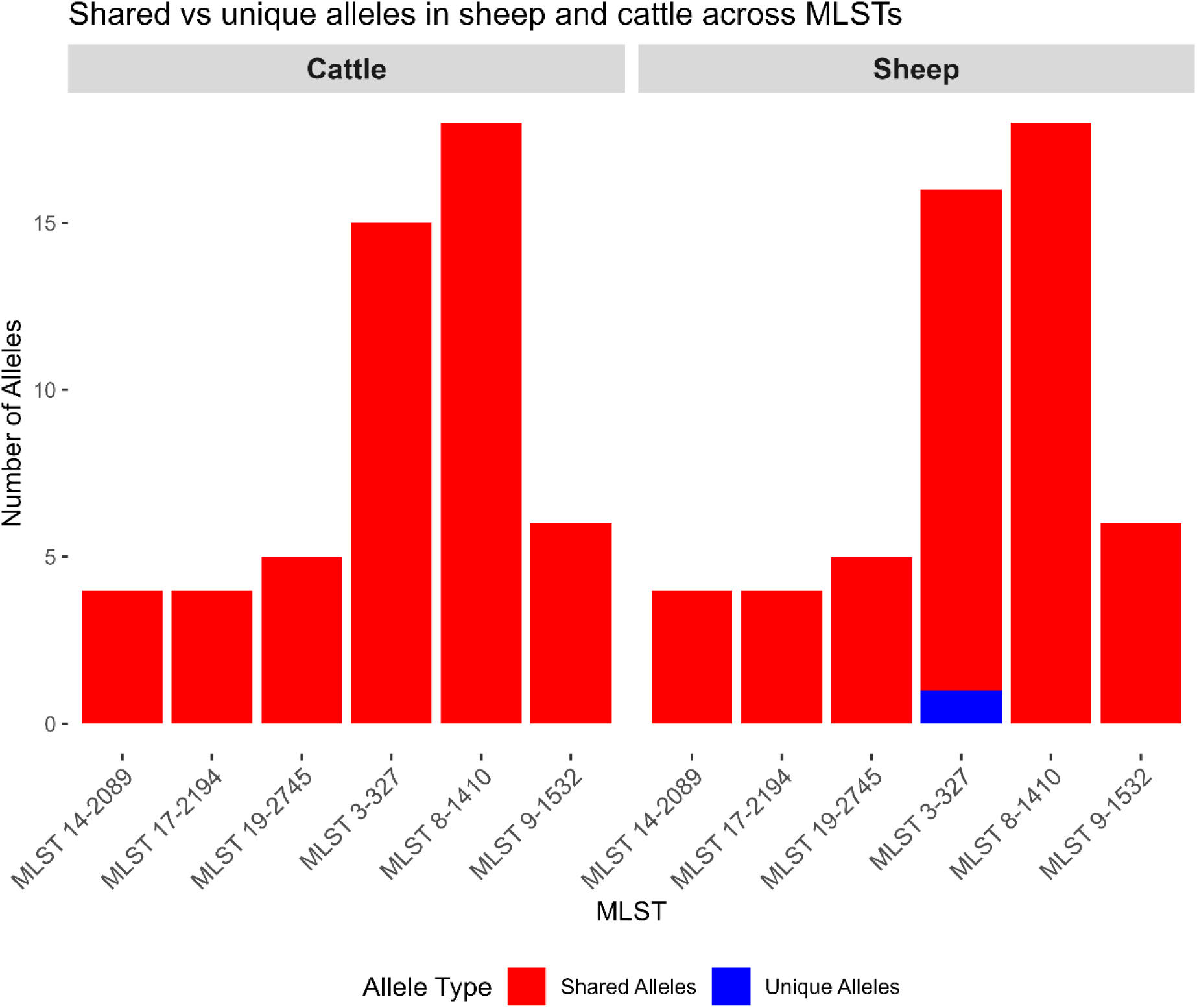
Unique and shared alleles observed in each MLST from all in cattle and sheep-derived *F. hepatica* populations

**Table 4.** Summary of alleles and the single nucleotide polymorphisms (SNPs) found at 6 MLST marker loci in 15 cattle and 27 sheep-derived *F. hepatica* populations from natural infections in the UK.

| MLST marker | Host | Alleles | Total SNPs in all alleles | SNPs range per allele | Average SNPs per Allele | s.d. SNPs per Allele | Host | Alleles | Total SNPs in all alleles | SNPs range per allele | Average SNPs per Allele | s.d. SNPs per Allele |
| --- | --- | --- | --- | --- | --- | --- | --- | --- | --- | --- | --- | --- |
| 3-327 | Cattle | 15 | 59 | 1 – 8 | 3.93 | 2.63 | Sheep | 16 | 63 | 1 – 8 | 3.94 | 2.54 |
| 8-1410 | Cattle | 18 | 37 | 1 – 4 | 2.06 | 0.87 | Sheep | 18 | 37 | 1 – 4 | 2.06 | 0.87 |
| 9-1532 | Cattle | 6 | 1 | 0 – 1 | 0.17 | 0.41 | Sheep | 6 | 1 | 0 – 1 | 0.17 | 0.41 |
| 14-2089 | Cattle | 4 | 6 | 0 - 3 | 1.5 | 1.73 | Sheep | 4 | 6 | 0 – 3 | 1.5 | 1.73 |
| 17-2194 | Cattle | 4 | 10 | 1 – 4 | 2.5 | 1.73 | Sheep | 4 | 10 | 1 – 4 | 2.5 | 1.73 |
| 19-2745 | Cattle | 5 | 12 | 0 – 9 | 2.4 | 3.78 | Sheep | 5 | 12 | 0 – 9 | 2.4 | 3.78 |

### Genetic diversity in MLSTs

Based on six MLST loci, expected heterozygosity (He) in *F. hepatica* showed considerable variation between loci, with overall moderate mean genetic diversity in both cattle- and sheep-derived populations. At MLST 3-327, mean He (±SD) was lower in cattle (0.316 ± 0.217; range 0–0.571) than in sheep (0.441 ± 0.104; range 0.270–0.600). At MLST 8-1410, similar levels of diversity were observed in cattle (0.536 ± 0.059; range 0.400–0.643) and sheep (0.541 ± 0.039; range 0.433–0.611). Low diversity was recorded at MLST 9-1532 in both hosts (cattle 0.215 ± 0.214; range 0–0.50; sheep 0.242 ± 0.193; range 0–0.50). In contrast, higher variability was observed at MLST 14-2089 (cattle 0.292 ± 0.377; range 0–1; sheep 0.492 ± 0.259; range 0–1), MLST 17-2194 (cattle 0.700 ± 0.399; range 0–1; sheep 0.698 ± 0.277; range 0–1), and MLST 19-2745 (cattle 0.370 ± 0.351; range 0–0.667; sheep 0.450 ± 0.371; range 0–1) Supplementary Figure 3.

### Population sub-structure analysis

When comparing cattle and sheep populations, mean F_ST_ values for five MLST markers showed negative values (–0.06552 to –0.34577), indicating largely no genetic differentiation between cattle- and sheep-derived *F. hepatica* populations. Pairwise comparisons of F_ST_ values between the 15 cattle- and 27 sheep-derived *F. hepatica* populations, and AMOVA findings, further confirmed that most genetic differentiation was within populations rather than between them, demonstrating a lack of host-associated genetic sub-structure at these loci. Only a single marker, MLST 3-327, had a positive average F_ST_ value (0.08), suggesting limited genetic differentiation both within and between populations (Figure 6, Table 5). However, based on all six MLST markers, some populations showed significant genetic differentiation, for example, sheep-derived populations from North West England (P5S, P6S, P42S), and South West England (P27S), and a cattle-derived population from the Scottish Borders (P13C), from other cattle and sheep-derived populations (Supplementary Table 8, Supplementary Table 9).

**Figure 6.**
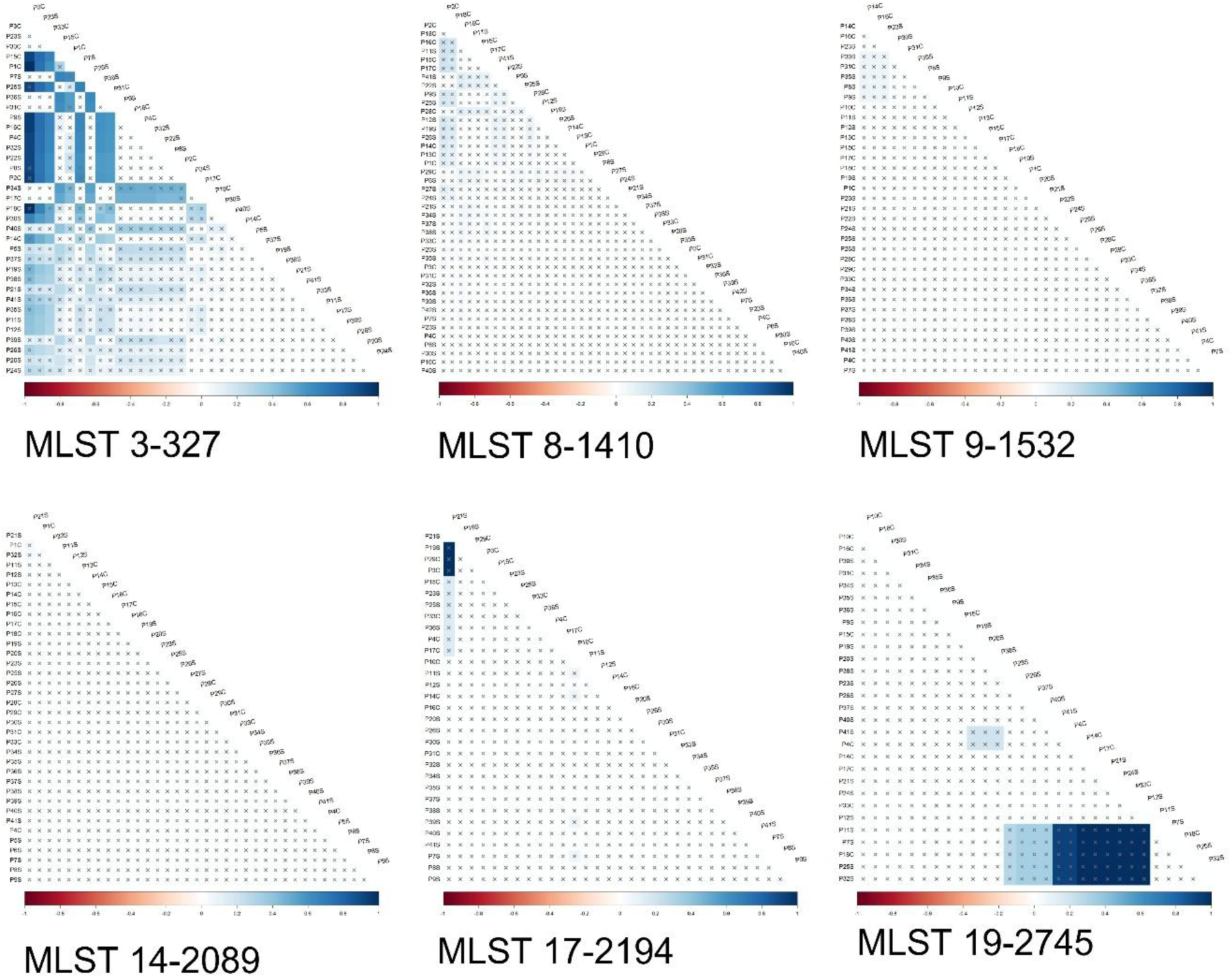
Pairwise comparison of F_ST_ values observed in 15 cattle and 27 sheep-derived *F. hepatica* populations from six MLSTs.

**Table 5.**
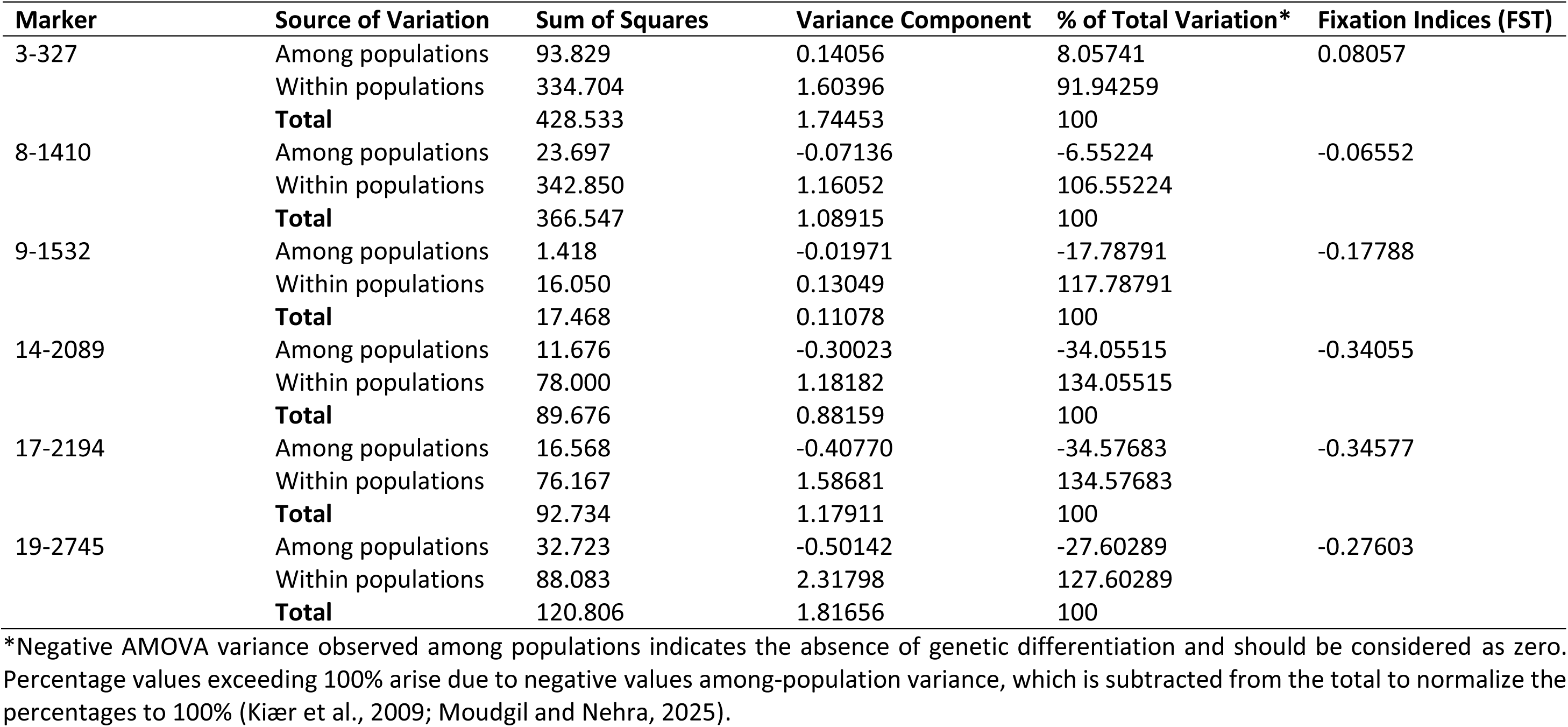
AMOVA results of 6 MLST markers in 27 sheep and 15 cattle-derived *F. hepatica* populations showing genetic variations between and within populations.

| Marker | Source of Variation | Sum of Squares | Variance Component | % of Total Variation* | Fixation Indices (FST) |
| --- | --- | --- | --- | --- | --- |
| 3-327 | Among populations | 93.829 | 0.14056 | 8.05741 | 0.08057 |
|  | Within populations | 334.704 | 1.60396 | 91.94259 |  |
|  | **Total** | 428.533 | 1.74453 | 100 |  |
| 8-1410 | Among populations | 23.697 | -0.07136 | -6.55224 | -0.06552 |
|  | Within populations | 342.850 | 1.16052 | 106.55224 |  |
|  | **Total** | 366.547 | 1.08915 | 100 |  |
| 9-1532 | Among populations | 1.418 | -0.01971 | -17.78791 | -0.17788 |
|  | Within populations | 16.050 | 0.13049 | 117.78791 |  |
|  | **Total** | 17.468 | 0.11078 | 100 |  |
| 14-2089 | Among populations | 11.676 | -0.30023 | -34.05515 | -0.34055 |
|  | Within populations | 78.000 | 1.18182 | 134.05515 |  |
|  | **Total** | 89.676 | 0.88159 | 100 |  |
| 17-2194 | Among populations | 16.568 | -0.40770 | -34.57683 | -0.34577 |
|  | Within populations | 76.167 | 1.58681 | 134.57683 |  |
|  | **Total** | 92.734 | 1.17911 | 100 |  |
| 19-2745 | Among populations | 32.723 | -0.50142 | -27.60289 | -0.27603 |
|  | Within populations | 88.083 | 2.31798 | 127.60289 |  |
|  | **Total** | 120.806 | 1.81656 | 100 |  |

\*Negative AMOVA variance observed among populations indicates the absence of genetic differentiation and should be considered as zero. Percentage values exceeding 100% arise due to negative values among-population variance, which is subtracted from the total to normalize the percentages to 100% (Kiær et al., 2009; Moudgil and Nehra, 2025).

PCoA plots were generated from distance matrices to assess the genetic distinctiveness of alleles from six MLSTs and overlapping clusters of identical alleles were observed between cattle- and sheep-derived populations from different regions of the UK (Figure 7). Thus, results from PCoA analysis reinforced previous findings and highlighted a lack of clear geographical or host-associated genetic sub-structuring.

**Figure 7.**
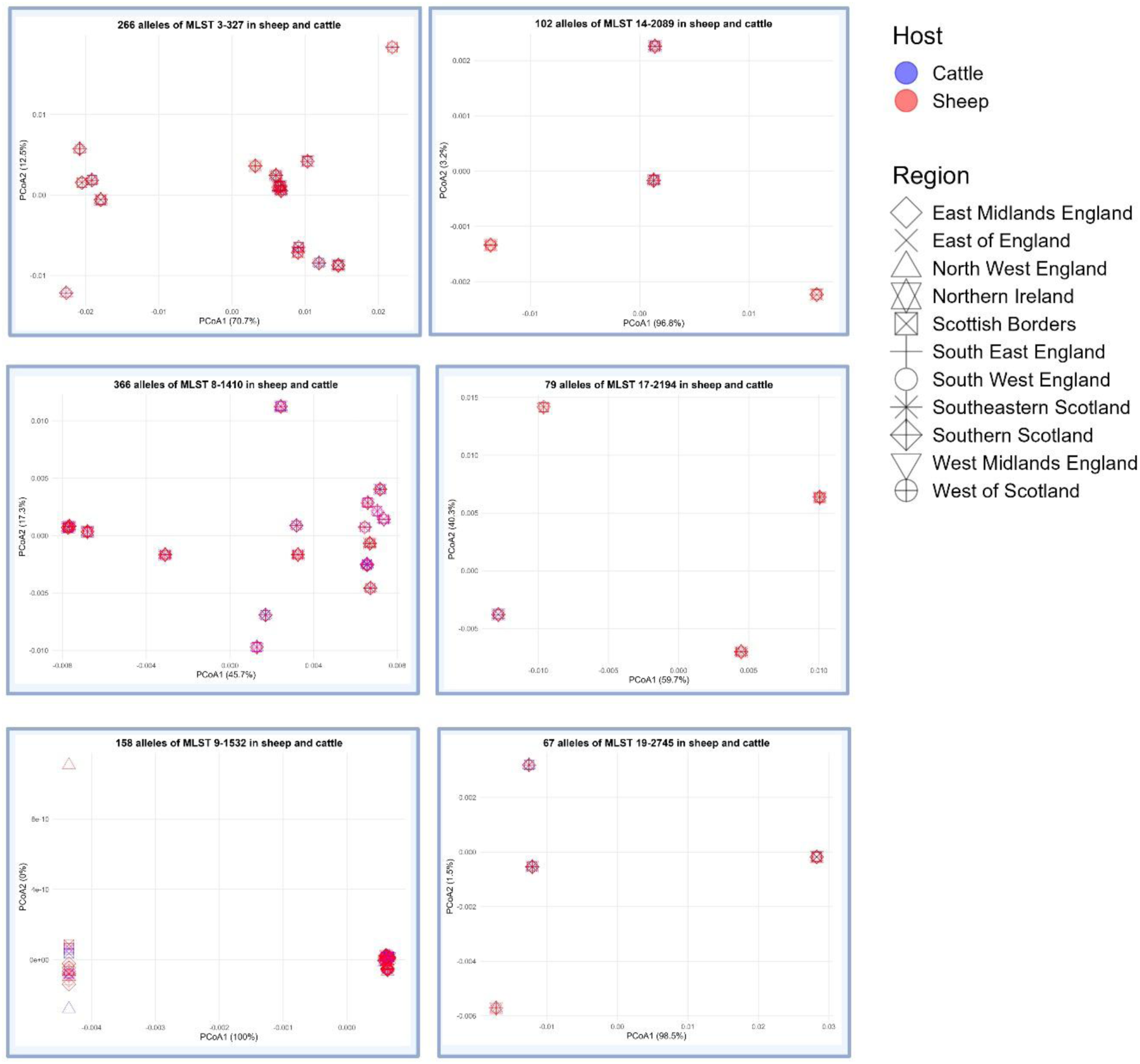
Principal coordinate analysis (PCoA) of alleles from six MLST markers in 15 cattle-and 27 sheep-derived *F. hepatica* populations across the UK. The plot illustrates the regional distribution of alleles using different shapes, sheep hosts are indicated in red and cattle in blue. The two principal coordinates (PCoA1 and PCoA2) explained the proportions of variation for each MLST marker including 266 alleles of MLST 3-327 (70.7 % and 12.5 %), 366 alleles of MLST 8-1410 (45.7 % and 17.3 %), 158 alleles of MLST 9-1532 (100 % and 00 %), 102 alleles of MLST 14-2089 (96.8 % and 3.2 %), 79 alleles of MLST 17-2194 (59.7 % and 40.3 %), and 67 alleles of MLST 19-2745 (98.5% and 1.5 %)

## Discussion

This study presents the design, testing and validation of a new panel of six MLST markers for population-genetic studies of *F. hepatica*, applicable to DNA extracted from adult worms isolated from livers and eggs purified from faeces. These MLST markers were mined from WGS data using two UK *F. hepatica* isolates and available reference genomes in the NCBI database and have been adapted for a next generation sequencing platform. On applying these makers to *F. hepatica* populations derived from cattle and sheep in the UK we observed overlapping allele patterns in sheep and cattle from different regions of the UK, indicating high gene flow in both hosts and limited genetic structuring between populations. The *F. hepatica* genome (1.2 to 1.5 Gb) is one of the largest among parasites and contains a notable duplication rate. In Dec 2014, the University of Liverpool, UK deposited the first *F. hepatica* genome assembly; however, its scaffold N50 quality was low N50=204 kb (BioProject PRJEB6687) (Cwiklinski et al., 2015b). Later in 2018, as technology advanced, assembly quality improved to N50 = 1.9 Mb (BioProject PRJEB25283), and in 2023, to N50 = 9.8 Mb (BioProject PRJEB58756). Additionally, a genome assembly (N50 = 161.1 kb) of *F. hepatica* (BioProject PRJNA179522) was documented from the USA (McNulty et al., 2017). We employed three reference genomes for mapping of our *F. hepatica* Illumina PE150 paired-end reads and found high alignment rates. The GC content of our WGS data was comparable to that of other available *F. hepatica* reference genomes (Cwiklinski et al., 2015b; McNulty et al., 2017). The duplication rate in the merged paired reads in our samples approximately 25% was lower than the previously reported genome-wide 32% duplication rate (Cwiklinski et al., 2015b). The overall alignment rates and mapping percentages of our genomic data showed high similarity to reference genomes of *F. hepatica* from the UK and the USA (Cwiklinski et al., 2015b; McNulty et al., 2017).

Hundreds of polymorphic fragments were mined from three available genomes for the construction of genomic markers. We used a threshold of ≥5 SNPs per genomic fragment to construct each MLST marker, which was higher than the threshold of two SNPs per genomic region used in a previous report on *C. parvum* (Troell et al., 2025). The successful amplification of 18 out of the 20 initial MLST markers demonstrated the feasibility of using SNP-rich fragments for MLST marker development. However, only six markers passed through all validation steps. Similarly, for *F. hepatica* microsatellite marker development, 35 loci were initially tested, and 15 were successfully optimised and multiplexed (Cwiklinski et al., 2015a).

The liver fluke samples used to validate the MLST markers were obtained from naturally infected sheep and cattle, collected through faecal examination and liver inspection across multiple regions of the UK (Abbas et al., 2026). Amplification success for MLSTs was higher in worm DNA than in *Fasciola* egg DNA. This is to be expected due to lower template concentration and the presence of inhibitors (Kikuchi et al., 2010). Six markers were selected that showed good amplification in both worm and egg DNA samples. The detection rates of these six MLST markers in both host species indicated that these loci can be suitable for population genetic studies.

The selected MLST markers showed limited allelic variation across the *F. hepatica* populations analysed; however, the total, average, and range of SNPs observed in alleles from natural infections indicated that MLSTs 3-327, 8-1410, 14-2089, 17-2194, and 19-2745 are polymorphic loci. Different studies using polymorphic microsatellite markers have reported considerable genetic variation in *F. hepatica* isolates from the UK (Beesley et al., 2017), Argentina (Beesley et al., 2021), and South Africa (Nukeri et al., 2025). Geographical patterns of genetic diversity have been reported in liver fluke populations using ribosomal and mitochondrial markers in Eastern Europe (Teofanova et al., 2011), and in China (Tang et al., 2021) and using ribosomal, mitochondrial and microsatellite markers in German dairy cattle (Hecker et al., 2025). Therefore, using MLST markers on a larger sample from different geographic areas, along with other markers, is a good option for studying population genetics.

Moreover, the presence of monomorphic loci with a single SNP in several populations at MLST 9-1532 suggests that some genomic regions are highly conserved, whereas other loci, such as 3-327, 8-1410, 14-2089, 17-2194 and 19-2745 show higher levels of polymorphism. Differences in polymorphism across loci can be due to variation in mutation rates at the genomic level. Some loci may reside in regions with higher mutation rates, leading to greater allelic variation, whereas conserved genomic regions exhibit lower variability. Additionally, mutations at different loci may influence the genetic diversity. Microsatellites generally exhibit higher allelic variability due to their higher mutation rates, whereas SNP-based genome-wide markers provide unbiased estimates of genetic diversity (Fischer et al., 2017).

In this study, high allelic richness was observed in *F. hepatica* populations in the Scottish Borders, Southern Scotland, and parts of South-West England, in line with previous mitochondrial and nuclear microsatellite DNA studies reporting high allelic richness in the UK and Ireland (Beesley et al., 2017; Walker et al., 2007). Parasite populations from the Scottish Borders and South-West England from cattle and sheep hosts, respectively, showed genetic differentiation, highlighting the potential mixing of metacercariae on pasture or variation in sequence read depth across loci among different parasite populations. However, this needs further investigation on a larger sample size populations from both hosts.

Few sheep-derived populations from North-West England exhibited genetic differentiation, although they had low allelic variation, and multiple markers yielded few or no alleles. It has been reported that North-West England is an endemic area for liver fluke, with a prevalence of 85.1% in milk samples from high-yielding dairy herds (Howell et al., 2015). However, the low number of allelic variation patterns and genetic differentiation observed in our work may be due to a higher degree of clonal transmission and limited mixing of the metacercariae of the parasite (Beesley et al., 2017; Vilas et al., 2012).

Sheep-derived populations showed higher allelic variation than cattle-derived populations, however, heterozygosity was in moderate levels in cattle and sheep derived *F. hepatica* populations. In our study, the number of parasite populations in sheep was about twice that in cattle, and sequence read depth also varied in different populations and at different MLST loci which may have influenced the allelic variations. In northwestern Spain, analysis of eight allozymic and four microsatellite markers across 20 infra populations derived from 10 cattle and 10 sheep showed contrasting patterns of population genetic structure between hosts. Sheep-derived populations showed a non-random occurrence of clones, greater population structuring, and higher levels of gene diversity compared with cattle-derived populations. (Vilas et al., 2012).

Our findings indicate that most genetic differentiation occurs within rather than between cattle- and sheep-derived populations, which may facilitate parasitic adaptation to host and environmental conditions (Cwiklinski et al., 2015b). No genetic differences was found between *F. hepatica* populations from three regions of China, however, genetic differentiation was observed within sheep derived populations using mitochondrial markers (Xifeng et al., 2022). Further, identical alleles were recorded in sheep and cattle, indicating a lack of host-associated genetic sub-structuring in *F. hepatica*. The multiple occurrences of similar alleles in populations and hosts which we observed indicated high gene flow in both hosts. High gene flow could facilitate the spread of resistant alleles once they arise. Indeed, Beesley et al., (2017) proposed that high gene flow and limited population structure could allow rapid dissemination of resistance. Cross-species transmission of clonemates can occur through grazing in the same pastures (Beesley et al., 2021). Other possibilities include animal sales, livestock movement between farms (Walker et al., 2007), and snail transfer by wildlife, humans, or livestock (Mas-Coma et al., 2009). Previously, Beesley et al. (2017) sampled over 1,500 adult flukes from cattle and sheep in the UK and found clonal parasites with similar multilocus genotypes in 61% of hosts. However, 84% of parasites had unique multilocus genotypes which is showing high genetic diversity within parasite populationsand estimated a selfing rate of no more than 2% and this diversity can be due to cross fertilisation. In contrast, in Spain, based on microsatellites and mitochondrial markers, contrasting genetic structures within cattle- and sheep-origin parasites has also been reported with more structured genetic variation in sheep derived infrapopulation (Vilas et al., 2012).

The present study has limitations. Firstly, we used pooled DNA samples to test MLST markers, which limits us to estimating Hardy–Weinberg equilibrium, calculating inbreeding coefficients (F_IS_), and probability of sexual origin (Psex). Biallelic data are required to calculate these parameters, which are important for assessing self-fertilisation and cross-fertilisation in *F. hepatica* populations. Secondly, the population sizes were small and uneven across host groups, cattle (n = 15) and sheep (n = 27), which can introduce biases in estimates of allelic diversity between cattle- and sheep-derived parasite populations. Thirdly, we cannot find gene annotation data in the database for these MLST genomic regions, which restricts us from analysing the functional interpretation of allelic variation. Next, MLST markers are based solely on SNPs and are extracted from different genome scaffolds, which may evolve more slowly than microsatellites. Finally, a significant limitation of this study was the substantial variation in sequencing read depth among MLST markers, likely resulting from differences in amplification efficiency between loci. Consequently, the number of alleles identified varied considerably across markers. Importantly, a significant positive association was detected between sequencing read counts and the number of alleles observed per locus, indicating that loci with higher read depth generated a higher number of alleles. Therefore, the variations in diversity may reflect technical variation associated with sequencing coverage rather than true biological diversity, and this should be considered when interpreting patterns of population genetic structure.

Given these limitations, future studies need to use individual parasitic worm-based sequencing with larger, balanced sampling from both host and parasite, and to calculate Hardy–Weinberg equilibrium, inbreeding coefficients (F_IS_), and Psex (Beesley et al., 2021). Further, comparisons of microsatellite marker systems with MLST markers reported in this study are highly recommended to refine our understanding of *F. hepatica* population genetics. Moreover, collection of environmental and farm-management data alongside samples can help identify potential drivers of local genetic population structure.

In conclusion, we designed, developed, and validated a panel of six genome-wide MLST markers for population genetic analysis of *F. hepatica* and demonstrated that the markers functioned in both adult worms and egg DNA samples. Geographic variation in allelic diversity was observed, with the highest diversity in the Scottish Borders, Southern Scotland, and South-West England, while parasite populations from the North-West of England displayed low allelic richness. Sheep- and cattle-derived populations showed gene flow between the two host species, as alleles were shared in both hosts. One MLST marker was monomorphic, and five markers showed polymorphism within the study. This highlights the importance of multilocus approaches in population genetics. Overall, the results showed the potential of MLST markers for exploring regional and host-associated population genetics of *F. hepatica* populations and provides an additional tool for future evolutionary studies.

## Ethical statement

Non-invasive collection of faecal samples was approved by the NASPA (Non-Animal Scientific Procedures Act) sub-committee of AWERB, University of Surrey, UK, under the reference NASPA-2122-04 for the project “Developing Novel Rapid Diagnostics for Neglected Parasitic Diseases.” Adult flukes were collected at licensed slaughterhouses and through post-mortem examination. Completion of a University of Surrey SAGE-AR (ID 638929-638920-101535552) indicated that no formal ethical approval was required for adult fluke sampling.

## Supplementary information

Supplementary Figure 1 to 3

Supplementary Tables 1 to 9

## CRediT authorship contribution statement

Muhammad Abbas: conceptualisation, investigation, methodology, bioinformatics, validation, visualisation, data curations and analysis, writing original draft, review and editing; Kezia Kozel: resources, review and editing; Olukayode Daramola: review and editing, supervision; Nick Selemetas: review and editing, supervision; Mark W. Robinson: review and editing, resources; Eric R Morgan: conceptualisation, funding acquisition, supervision, review and editing; Umer Chaudhry: conceptualisation, review and editing, supervision; Martha Betson: conceptualisation, review and editing, supervision, funding acquisition, project administration.

## Acknowledgements

Part of this work was carried out using the computational HPC facilities and support provided by the Research Computing Services team within IT Services at the University of Surrey, specifically the Eureka2 HPC cluster (https://docs.pages.surrey.ac.uk/research_computing/hpc/clusters/eureka2.html).

This research was funded in whole, or in part, by the UK Research and Innovation (UKRI), Biotechnology and Biological Sciences Research Council (BBSRC) through the FoodBioSystems Doctoral Training Programme (BB/T008776/1) and the Sir Halley Stewart Trust (3153). For Open Access, the authors have applied a Creative Commons Attribution (CC BY) public copyright licence to any Author Accepted Manuscript version arising from this submission. We sincerely acknowledge all farmers and registered veterinary practitioners in the UK and especially Dr. Iñaki Deza-Cruz (The Royal (Dick) School of Veterinary Studies and The Roslin Institute, The University of Edinburgh, Easter Bush Veterinary Centre, Midlothian, EH25 9RG), and Dr. Sai Fingerhood (Department of Veterinary Pathology, University of Nottingham, UK), for their valuable assistance in sample collection.

## Rights and permissions

All sequencing data reported in the paper are available under NCBI BioProject ID PRJNA1431397 and accession numbers SAMN56308754 to SAMN56308884; SAMN56308979 to SAMN56309022; SAMN56321787 and SAMN56321788.

In addition, sequence data, script, and codes are available at the Mendeley database https://data.mendeley.com/datasets/gc8fwkxdhy/1 and https://github.com/drmuhammadabbas810/New-tools-for-sustainable-control-of-liver-fluke-in-ruminants

All other data are reported in the paper and associated supplementary material.

## References

Abbas, M., Kozel, K., Daramola, O., Selemetas, N., Ali, Q., Ashraf, S., Ibrahim, I., Deza-Cruz, I., Fingerhood, S., Robinson, M.W., Morgan, E.R., Chaudhry, U., Betson, M., 2026. Development of a qPCR assay for *Fasciola* spp. identification and deep amplicon sequencing method for differentiation of fluke species in UK livestock. PLoS Negl. Trop. Dis. 20, e0014006. 10.1371/journal.pntd.0014006

Ballard, J.W.O., Whitlock, M.C., 2004. The incomplete natural history of mitochondria. Mol. Ecol. 13, 729–744. 10.1046/j.1365-294X.2003.02063.x

Bathke, J., Lühken, G., 2021. OVarFlow: a resource optimized GATK 4 based Open source Variant calling workFlow. BMC Bioinformatics 22, 402. 10.1186/s12859-021-04317-y

Beesley, N.J., Attree, E., Vázquez-Prieto, S., Vilas, R., Paniagua, E., Ubeira, F.M., Jensen, O., Pruzzo, C., Álvarez, J.D., Malandrini, J.B., Solana, H., Hodgkinson, J.E., 2021. Evidence of population structuring following population genetic analyses of *Fasciola hepatica* from Argentina. Int. J. Parasitol. 51, 471–480. 10.1016/j.ijpara.2020.11.007

Beesley, N.J., Williams, D.J.L., Paterson, S., Hodgkinson, J., 2017. *Fasciola hepatica* demonstrates high levels of genetic diversity, a lack of population structure and high gene flow: possible implications for drug resistance. Int. J. Parasitol. 47, 11–20. 10.1016/j.ijpara.2016.09.007

Bodenhofer, U., Bonatesta, E., Horejš-Kainrath, C., Hochreiter, S., 2015. msa: an R package for multiple sequence alignment. Bioinformatics 31, 3997–3999. 10.1093/bioinformatics/btv494

Bogenhagen, D., Clayton, D.A., 1974. The number of mitochondrial deoxyribonucleic acid genomes in mouse L and human HeLa cells: quantitative isolation of mitochondrial deoxyribonucleaic acid. J. Biol. Chem. 249, 7991–7995. 10.1016/S0021-9258(19)42063-2

Van den Brand, T., 2025. ggh4x: Hacks for “ggplot2.” https://cran.r-project.org/web/packages/ggh4x/index.html

Calvani, N.E.D., Šlapeta, J., 2021. *Fasciola* species introgression: just a fluke or something more? Trends Parasitol. 37, 25–34. 10.1016/j.pt.2020.09.008

Charif, D., Lobry, J.R., 2007. SeqinR 1.0-2: A Contributed Package to the R Project for Statistical Computing Devoted to Biological Sequences Retrieval and Analysis, in: Bastolla, U., Porto, M., Roman, H.E., Vendruscolo, M. (Eds.), Structural Approaches to Sequence Evolution: Molecules, Networks, Populations. Springer, Berlin, Heidelberg, pp. 207–232. 10.1007/978-3-540-35306-5_10

Charlier, J., van der Voort, M., Kenyon, F., Skuce, P., Vercruysse, J., 2014. Chasing helminths and their economic impact on farmed ruminants. Trends Parasitol. 30, 361–367. 10.1016/j.pt.2014.04.009

Chen, S., Zhou, Y., Chen, Y., Gu, J., 2018. fastp: an ultra-fast all-in-one FASTQ preprocessor. Bioinformatics 34, i884–i890. 10.1093/bioinformatics/bty560

Criscione, C.D., Blouin, M.S., 2006. Minimal selfing, few clones, and no among-host genetic structure in a hermaphroditic parasite with asexual larval propagation. Evolution 60, 553–562. 10.1554/05-421.1

Cwiklinski, K., Allen, K., LaCourse, J., Williams, D.J., Paterson, S., Hodgkinson, J.E., 2015a. Characterisation of a novel panel of polymorphic microsatellite loci for the liver fluke, *Fasciola hepatica*, using a next generation sequencing approach. Infect. Genet. Evol. 32, 298–304. 10.1016/j.meegid.2015.03.014

Cwiklinski, K., Dalton, J.P., Dufresne, P.J., La Course, J., Williams, D.J., Hodgkinson, J., Paterson, S., 2015b. The *Fasciola hepatica* genome: gene duplication and polymorphism reveals adaptation to the host environment and the capacity for rapid evolution. Genome Biol. 16, 71. 10.1186/s13059-015-0632-2

Dalton, J.P., 2021. Fasciolosis. CABI.

Diosque, P., Tomasini, N., Lauthier, J.J., Messenger, L.A., Rumi, M.M.M., Ragone, P.G., Alberti-D’Amato, A.M., Brandán, C.P., Barnabé, C., Tibayrenc, M., Lewis, M.D., Llewellyn, M.S., Miles, M.A., Yeo, M., 2014. Optimized multilocus sequence typing (MLST) scheme for *Trypanosoma cruzi*. PLoS Negl. Trop. Dis. 8, e3117. 10.1371/journal.pntd.0003117

Dreyfuss, G., Rondelaud, D., 1994. *Fasciola hepatica*: a study of the shedding of cercariae from *Lymnaea truncatula* raised under constant conditions of temperature and photoperiod. Parasite Paris Fr. 1, 401–404. 10.1051/parasite/1994014401

Elliott, T., Muller, A., Brockwell, Y., Murphy, N., Grillo, V., Toet, H.M., Anderson, G., Sangster, N., Spithill, T.W., 2014. Evidence for high genetic diversity of NAD1 and COX1 mitochondrial haplotypes among triclabendazole resistant and susceptible populations and field isolates of *Fasciola hepatica* (liver fluke) in Australia. Vet. Parasitol. 200, 90–96. 10.1016/j.vetpar.2013.11.019

Excoffier, L., Lischer, H.E.L., 2010. Arlequin suite ver 3.5: a new series of programs to perform population genetics analyses under Linux and Windows. Mol. Ecol. Resour. 10, 564–567. 10.1111/j.1755-0998.2010.02847.x

Fischer, M.C., Rellstab, C., Leuzinger, M., Roumet, M., Gugerli, F., Shimizu, K.K., Holderegger, R., Widmer, A., 2017. Estimating genomic diversity and population differentiation – an empirical comparison of microsatellite and SNP variation in Arabidopsis halleri. BMC Genomics 18, 69. 10.1186/s12864-016-3459-7

Fletcher, H.L., Hoey, E.M., Orr, N., Trudgett, A., Fairweather, I., Robinson, M.W., 2004. The occurrence and significance of triploidy in the liver fluke, *Fasciola hepatica*. Parasitology 128, 69–72. 10.1017/S003118200300427X

Galtier, N., Nabholz, B., Glémin, S., Hurst, G.D.D., 2009. Mitochondrial DNA as a marker of molecular diversity: a reappraisal. Mol. Ecol. 18, 4541–4550. 10.1111/j.1365-294X.2009.04380.x

Hecker, A.S., Raulf, M.-K., König, S., May, K., Strube, C., 2025. Population genetic analysis of the liver fluke *Fasciola hepatica* in German dairy cattle reveals high genetic diversity and associations with fluke size. Parasit. Vectors 18, 51. 10.1186/s13071-025-06701-6

Hodel, R.G.J., Segovia-Salcedo, M.C., Landis, J.B., Crowl, A.A., Sun, M., Liu, X., Gitzendanner, M.A., Douglas, N.A., Germain-Aubrey, C.C., Chen, S., Soltis, D.E., Soltis, P.S., 2016. The report of my death was an exaggeration: A review for researchers using microsatellites in the 21st century1. Appl. Plant Sci. 4, apps.1600025. 10.3732/apps.1600025

Hodgkinson, J.E., Cwiklinski, K., Beesley, N., Hartley, C., Allen, K., Williams, D.J.L., 2018. Clonal amplification of *Fasciola hepatica* in *Galba truncatula*: within and between isolate variation of triclabendazole-susceptible and -resistant clones. Parasit. Vectors 11, 363. 10.1186/s13071-018-2952-z

Hoffman, J.I., Amos, W., 2005. Microsatellite genotyping errors: detection approaches, common sources and consequences for paternal exclusion. Mol. Ecol. 14, 599–612. 10.1111/j.1365-294X.2004.02419.x

Howell, A., Baylis, M., Smith, R., Pinchbeck, G., Williams, D., 2015. Epidemiology and impact of *Fasciola hepatica* exposure in high-yielding dairy herds. Prev Vet Med 121, 41–48. 10.1016/j.prevetmed.2015.05.013

Hurtrez-Boussès, S., Durand, P., Jabbour-Zahab, R., Guégan, J.-F., Meunier, C., Bargues, M.-D., Mas-Coma, S., Renaud, F., 2004. Isolation and characterization of microsatellite markers in the liver fluke (*Fasciola hepatica*). Mol. Ecol. Notes 4, 689–690. 10.1111/j.1471-8286.2004.00786.x

Jakobsson, M., Edge, M.D., Rosenberg, N.A., 2013. The relationship between F(ST) and the frequency of the most frequent allele. Genetics 193, 515–528. 10.1534/genetics.112.144758

John, B.C., Davies, D.R., Howell, A.K., Williams Diana J.L., Hodgkinson, J.E., 2020. Anaerobic fermentation results in loss of viability of *Fasciola hepatica* metacercariae in grass silage. Vet. Parasitol. 285, 109218. 10.1016/j.vetpar.2020.109218

Kiær, L.P., Felber, F., Flavell, A., Guadagnuolo, R., Guiatti, D., Hauser, T.P., Olivieri, A.M., Scotti, I., Syed, N., Vischi, M., van de Wiel, C., Jørgensen, R.B., 2009. Spontaneous gene flow and population structure in wild and cultivated chicory, Cichorium intybus L. Genet. Resour. Crop Evol. 56, 405–419. 10.1007/s10722-008-9375-1

Kikuchi, A., Sawamura, T., Kawase, N., Kitajima, Y., Yoshida, T., Daimaru, O., Nakakita, T., Itoh, S., 2010. Utility of Spermidine in PCR Amplification of Stool Samples. Biochem. Genet. 48, 428–432. 10.1007/s10528-009-9326-3

Kolde, R., 2025. pheatmap: Pretty Heatmaps. https://cran.r-project.org/web/packages/pheatmap/index.html

Ladoukakis, E.D., Zouros, E., 2017. Evolution and inheritance of animal mitochondrial DNA: rules and exceptions. J. Biol. Res. 24, 2. 10.1186/s40709-017-0060-4

Langmead, B., Salzberg, S.L., 2012. Fast gapped-read alignment with Bowtie 2. Nat. Methods 9, 357–359. 10.1038/nmeth.1923

Langmead, B., Wilks, C., Antonescu, V., Charles, R., 2019. Scaling read aligners to hundreds of threads on general-purpose processors. Bioinformatics 35, 421–432. 10.1093/bioinformatics/bty648

Lerch, A., Koepfli, C., Hofmann, N.E., Messerli, C., Wilcox, S., Kattenberg, J.H., Betuela, I., O’Connor, L., Mueller, I., Felger, I., 2017. Development of amplicon deep sequencing markers and data analysis pipeline for genotyping multi-clonal malaria infections. BMC Genomics 18, 864. 10.1186/s12864-017-4260-y

Mas-Coma, S., Bargues, M.D., Valero, M.A., 2018. Human fascioliasis infection sources, their diversity, incidence factors, analytical methods and prevention measures. Parasitology 145, 1665–1699. 10.1017/S0031182018000914

Mas-Coma, S., Valero, M.A., Bargues, M.D., 2009. Chapter 2 Fasciola, Lymnaeids and Human Fascioliasis, with a Global Overview on Disease Transmission, Epidemiology, Evolutionary Genetics, Molecular Epidemiology and Control, in: Advances in Parasitology. Academic Press, pp. 41–146. 10.1016/S0065-308X(09)69002-3

McKenna, A., Hanna, M., Banks, E., Sivachenko, A., Cibulskis, K., Kernytsky, A., Garimella, K., Altshuler, D., Gabriel, S., Daly, M., DePristo, M.A., 2010. The Genome Analysis Toolkit: a MapReduce framework for analyzing next-generation DNA sequencing data. Genome Res. 20, 1297–1303. 10.1101/gr.107524.110

McNulty, S.N., Tort, J.F., Rinaldi, G., Fischer, K., Rosa, B.A., Smircich, P., Fontenla, S., Choi, Y.-J., Tyagi, R., Hallsworth-Pepin, K., Mann, V.H., Kammili, L., Latham, P.S., Dell’Oca, N., Dominguez, F., Carmona, C., Fischer, P.U., Brindley, P.J., Mitreva, M., 2017. Genomes of *Fasciola hepatica* from the Americas reveal colonization with *Neorickettsia* endobacteria related to the agents of potomac horse and human sennetsu fevers. PLoS Genet. 13, e1006537. 10.1371/journal.pgen.1006537

Morgan, M., Pages, H., Obenchain, V., Hayden, N., 2016. Rsamtools: Binary alignment (BAM), FASTA, variant call (BCF), and tabix file import. R Package Version 1, 677–689.

Moudgil, A.D., Nehra, A.K., 2025. Mitochondrial genetic markers based phylogenetic analyses of Hyalomma dromedarii Koch, 1844 (Acari: Ixodidae). J. Genet. Eng. Biotechnol. 23, 100460. 10.1016/j.jgeb.2025.100460

Novobilský, A., Engström, A., Sollenberg, S., Gustafsson, K., Morrison, D.A., Höglund, J., 2014. Transmission patterns of *Fasciola hepatica* to ruminants in Sweden. Vet. Parasitol. 203, 276–286. 10.1016/j.vetpar.2014.04.015

Nukeri, S., Malatji, M.P., Mnisi, C.M., Chaisi, M., Mukaratirwa, S., 2025. Pilot study on the population genetics structure of *Fasciola hepatica* from seven provinces of South Africa. Front. Vet. Sci. 12. 10.3389/fvets.2025.1659523

Ooms, J., credetails, J.M. (Author of libxlsxwriter (see A. and C. files for details)) writexl author, 2025. writexl: Export Data Frames to Excel “xlsx” Format. https://cran.r-project.org/web/packages/writexl/index.html

Pagès, H., Aboyoun, P., Gentleman, R., DebRoy, S., 2025. Biostrings: Efficient manipulation of biological strings. R package version 2.78.0. doi:10.18129/B9.bioc.Biostrings

Peng, Q., Vijaya Satya, R., Lewis, M., Randad, P., Wang, Y., 2015. Reducing amplification artifacts in high multiplex amplicon sequencing by using molecular barcodes. BMC Genomics 16, 589. 10.1186/s12864-015-1806-8

Rehman, Z.U., Zahid, O., Rashid, I., Ali, Q., Akbar, M.H., Oneeb, M., Shehzad, W., Ashraf, K., Sargison, N.D., Chaudhry, U., 2020. Genetic diversity and multiplicity of infection in *Fasciola gigantica* isolates of Pakistani livestock. Parasitol. Int. 76, 102071. 10.1016/j.parint.2020.102071

Revell, L.J., 2024. phytools 2.0: an updated R ecosystem for phylogenetic comparative methods (and other things). PeerJ 12, e16505. 10.7717/peerj.16505

Sanz, C.R., Carbonell, J.D., Miró, G., Meana, A., 2025. Equine fasciolosis due to *Fasciola hepatica* in the Community of Madrid (Spain): First report of a rare parasitic infection in horses. Vet. Rec. Case Rep. 13, e70151. 10.1002/vrc2.70151

Sargison, N.D., Shahzad, K., Mazeri, S., Chaudhry, U., 2019. A high throughput deep amplicon sequencing method to show the emergence and spread of *Calicophoron daubneyi* rumen fluke infection in United Kingdom cattle herds. Vet. Parasitol. 268, 9–15. 10.1016/j.vetpar.2019.02.007

Schloss, P., Sl Westcott, Ryabin T, Hall Jr, M Hartmann, Hollister Eb, Lesniewski Ra, Oakley Bb, Parks Dh, Robinson Cj, Sahl Jw, Stres B, Thallinger Gg, Van Horn Dj, Weber Cf, 2009. Introducing mothur: open-source, platform-independent, community-supported software for describing and comparing microbial communities. Appl. Environ. Microbiol. 75. 10.1128/AEM.01541-09

Schwabl, P., Sánchez, J.M., Costales, J.A., Ocaña-Mayorga, S., Segovia, M., Carrasco, H.J., Hernández, C., Ramírez, J.D., Lewis, M.D., Grijalva, M.J., Llewellyn, M.S., 2020. Culture-free genome-wide locus sequence typing (GLST) provides new perspectives on *Trypanosoma cruzi* dispersal and infection complexity. PLOS Genet. 16, e1009170. 10.1371/journal.pgen.1009170

Schwartz, M., Vissing, J., 2002. Paternal inheritance of mitochondrial DNA. N. Engl. J. Med. 347, 576–580. 10.1056/NEJMoa020350

Semyenova, S.K., Morozova, E.V., Chrisanfova, G.G., Gorokhov, V.V., Arkhipov, I.A., Moskvin, A.S., Movsessyan, S.O., Ryskov, A.P., 2006. Genetic differentiation in eastern European and western Asian populations of the liver fluke, Fasciola hepatica, as revealed by mitochondrial NAD1 and COX1 genes. J. Parasitol. 92, 525–530. 10.1645/GE-673R.1

Šlapeta, J., Krücken, J., Rojas, A., Chambers, A., Melville, L.A., Martínez-Valladares, M., Canton, C., Francis, E.K., Zahid, O., Albuquerque, A.C.A., Bartley, D.J., Bassetto, C.C., Byrne, O., Colella, V., Costa-Junior, L.M., Doyle, S.R., Evans, M., Ghafar, A., Godoy, P., Hayashi, N., Helal, M.A., Huggins, L.G., Jabbar, A., Jones, R.A., Karani, B.E., Liron, J.P., Maté, L., McEvoy, A., Mohammedsalih, K.M., Mulcahy, G., Nielsen, M.K., Pafčo, B., Peachey, L.E., Robleto-Quesada, J., von Samson-Himmelstjerna, G., de Sousa-Paula, L.C., Gilleard, J.S., 2025. Ten simple rules for implementing deep amplicon sequencing in parasitology. Int. J. Parasitol. 10.1016/j.ijpara.2025.11.003

Snipen, L., Liland, K.H., 2025. microseq: basic biological sequence handling. https://cran.r-project.org/web/packages/microseq/index.html

Tang, M., Zhou, Y., Liu, Y., Cheng, N., Zhang, J., Xu, X., 2021. Molecular identification and genetic-polymorphism analysis of *Fasciola* flukes in Dali Prefecture, Yunnan Province, China. Parasitol. Int. 85, 102416. 10.1016/j.parint.2021.102416

Teofanova, D., Kantzoura, V., Walker, S., Radoslavov, G., Hristov, P., Theodoropoulos, G., Bankov, I., Trudgett, A., 2011. Genetic diversity of liver flukes (*Fasciola hepatica*) from Eastern Europe. Infect. Genet. Evol. 11, 109–115. 10.1016/j.meegid.2010.10.002

Troell, K., Stensvold, C.R., Sannella, A.R., Betson, M., Östlund, E., Chalmers, R.M., Chaudhry, U., Davidson, R., Davies, L., Ignatius, R., de Jong, A., Karadjian, G., Adjou, K., Klotz, C., Ptochos, S., Robinson, G., Roelfsema, J., Soba, B., Sroka, J., Vatta, P., Wensman, J.J., Cacciò, S.M., 2025. Design, development, and testing of a new multi-locus sequence typing scheme for the zoonotic pathogen *Cryptosporidium parvum*. Curr. Res. Parasitol. Vector-Borne Dis. 8, 100308. 10.1016/j.crpvbd.2025.100308

Utrera-Quintana, F., Covarrubias-Balderas, A., Olmedo-Juárez, A., Cruz-Aviña, J., Córdova-Izquierdo, A., Pérez-Mendoza, N., Villa-Mancera, A., 2022. Fasciolosis prevalence, risk factors and economic losses due to bovine liver condemnation in abattoirs in Mexico. Microb. Pathog. 173, 105851. 10.1016/j.micpath.2022.105851

Vázquez, A.A., Lounnas, M., Sánchez, J., Alba, A., Milesi, A., Hurtrez-Boussès, S., 2016. Genetic and infective diversity of the liver fluke *Fasciola hepatica* (Trematoda: Digenea) from Cuba. J. Helminthol. 90, 719–725. 10.1017/S0022149X15001029

Vilas, R., Vázquez-Prieto, S., Paniagua, E., 2012. Contrasting patterns of population genetic structure of *Fasciola hepatica* from cattle and sheep: implications for the evolution of anthelmintic resistance. Infect. Genet. Evol. J. Mol. Epidemiol. Evol. Genet. Infect. Dis. 12, 45–52. 10.1016/j.meegid.2011.10.010

Vyas, S., Mckay-Demeler, J., Ward, M., Calvani, N., 2026. A contemporary map of *Fasciola hepatica* distribution in sheep and cattle in New South Wales. Aust. Vet. J. 104, 25–36. 10.1111/avj.13465

Walker, S.M., Prodöhl, P.A., Fletcher, H.L., Hanna, R.E.B., Kantzoura, V., Hoey, E.M., Trudgett, A., 2007. Evidence for multiple mitochondrial lineages of *Fasciola hepatica* (liver fluke) within infrapopulations from cattle and sheep. Parasitol. Res. 101, 117–125. 10.1007/s00436-006-0440-4

Whitlock, M.C., 2011. G’ST and D do not replace FST. Mol. Ecol. 20, 1083–1091. 10.1111/j.1365-294X.2010.04996.x

Wickham, H., 2016. Data Analysis, in: Wickham, H. (Ed.), Ggplot2: Elegant Graphics for Data Analysis. Springer International Publishing, Cham, pp. 189–201. 10.1007/978-3-319-24277-4_9

Wickham, H., Averick, M., Bryan, J., Chang, W., McGowan, L., François, R., Grolemund, G., Hayes, A., Henry, L., Hester, J., Kuhn, M., Pedersen, T., Miller, E., Bache, S., Müller, K., Ooms, J., Robinson, D., Seidel, D., Spinu, V., Takahashi, K., Vaughan, D., Wilke, C., Woo, K., Yutani, H., 2019. Welcome to the tidyverse. J. Open Source Softw. 4, 1686. 10.21105/joss.01686

Wickham, H., Bryan, J., Posit, attribution), P. (Copyright holder of all R. code and all C. code without explicit copyright, code), M.K. (Author of included R., code), K.V. (Author of included libxls, code), C.L. (Author of included libxls, code), B.C. (Author of included libxls, code), D.H. (Author of included libxls, code), E.M. (Author of included libxls, 2025a. readxl: Read Excel Files. https://cran.r-project.org/web/packages/readxl/index.html

Wickham, H., François, R., Henry, L., Müller, K., Vaughan, D., Software, P., PBC, 2023. dplyr: A grammar of data manipulation. https://cran.r-project.org/web/packages/dplyr/index.html

Wickham, H., Henry, L., Software, P., PBC [cph, fnd, 2026a. purrr: Functional Programming Tools. https://cran.r-project.org/web/packages/purrr/index.html

Wickham, H., Hester, J., Francois, R., Bryan, J., Bearrows, S., Software, P., PBC [cph, fnd, library), https://github.com/mandreyel/ (mio, implementation), J.J. (grisu3, implementation), M.J. (grisu3, 2026b. readr: Read Rectangular Text Data. https://cran.r-project.org/web/packages/readr/index.html

Wickham, H., Vaughan, D., Girlich, M., Ushey, K., Software, P., PBC, 2025b. tidyr: Tidy Messy Data. https://cran.r-project.org/web/packages/tidyr/index.html

Wilke, C.O., 2025. cowplot: Streamlined Plot Theme and Plot Annotations for “ggplot2.” https://cran.r-project.org/web/packages/cowplot/index.html

Xifeng, W., Kai, Z., Guowu, Z., Zhiyuan, L., Yunxia, S., Chengcheng, N., Chunhui, J., Jun, Q., Qingling, M., Xuepeng, C., 2022. Molecular characteristics and genetic diversity of *Fasciola hepatica* from sheep in Xinjiang, China. J. Vet. Res. 66, 199–207. 10.2478/jvetres-2022-0018

Yasein, G., Zahid, O., Minter, E., Ashraf, K., Rashid, I., Shabbir, M.Z., Betson, M., Sargison, N.D., Chaudhry, U., 2022. A novel metabarcoded deep amplicon sequencing tool for disease surveillance and determining the species composition of *Trypanosoma* in cattle and other farm animals. Acta Trop. 230, 106416. 10.1016/j.actatropica.2022.106416

Zink, R.M., Barrowclough, G.F., 2008. Mitochondrial DNA under siege in avian phylogeography. Mol. Ecol. 17, 2107–2121. 10.1111/j.1365-294X.2008.03737.x

