## Supplementary Figure 1, 2 and 3 for "Development and validation of a multilocus sequence typing scheme for *Fasciola hepatica* using next-generation deep amplicon sequencing"

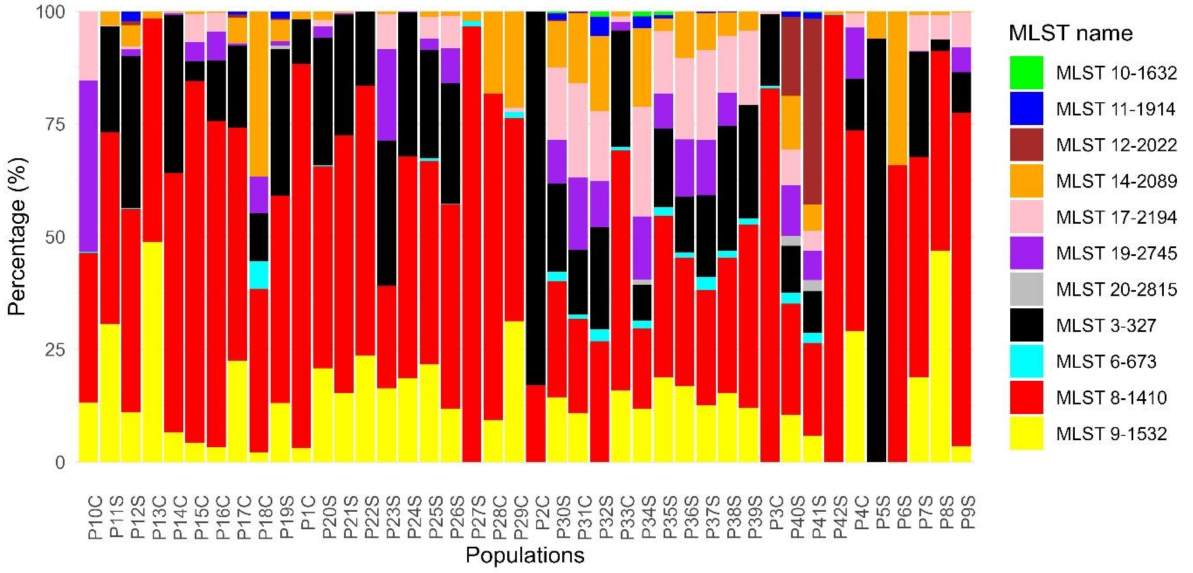


**Supplementary Fig. 1.** Distribution of sequence reads generated by MLST markers across different populations. The X-axis represents populations from cattle (C) and sheep (S), while the Y-axis shows the percentage of sequence reads from each MLST marker.

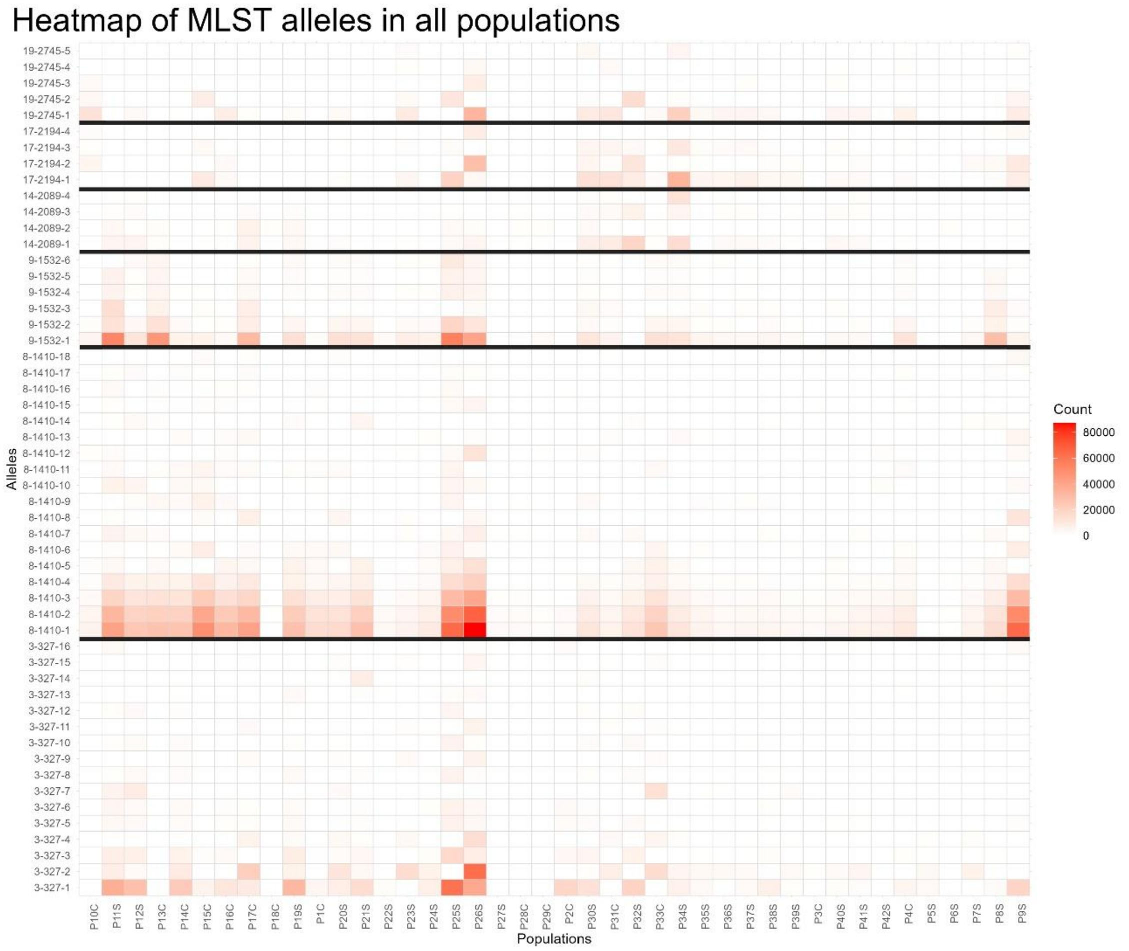


**Supplementary Fig. 2.** Heatmap of sequence reads of 53 alleles identified by 6 MLST markers in 27 sheep and 15 cattle-derived *F. hepatica* populations. A black line separates alleles for each MLST.

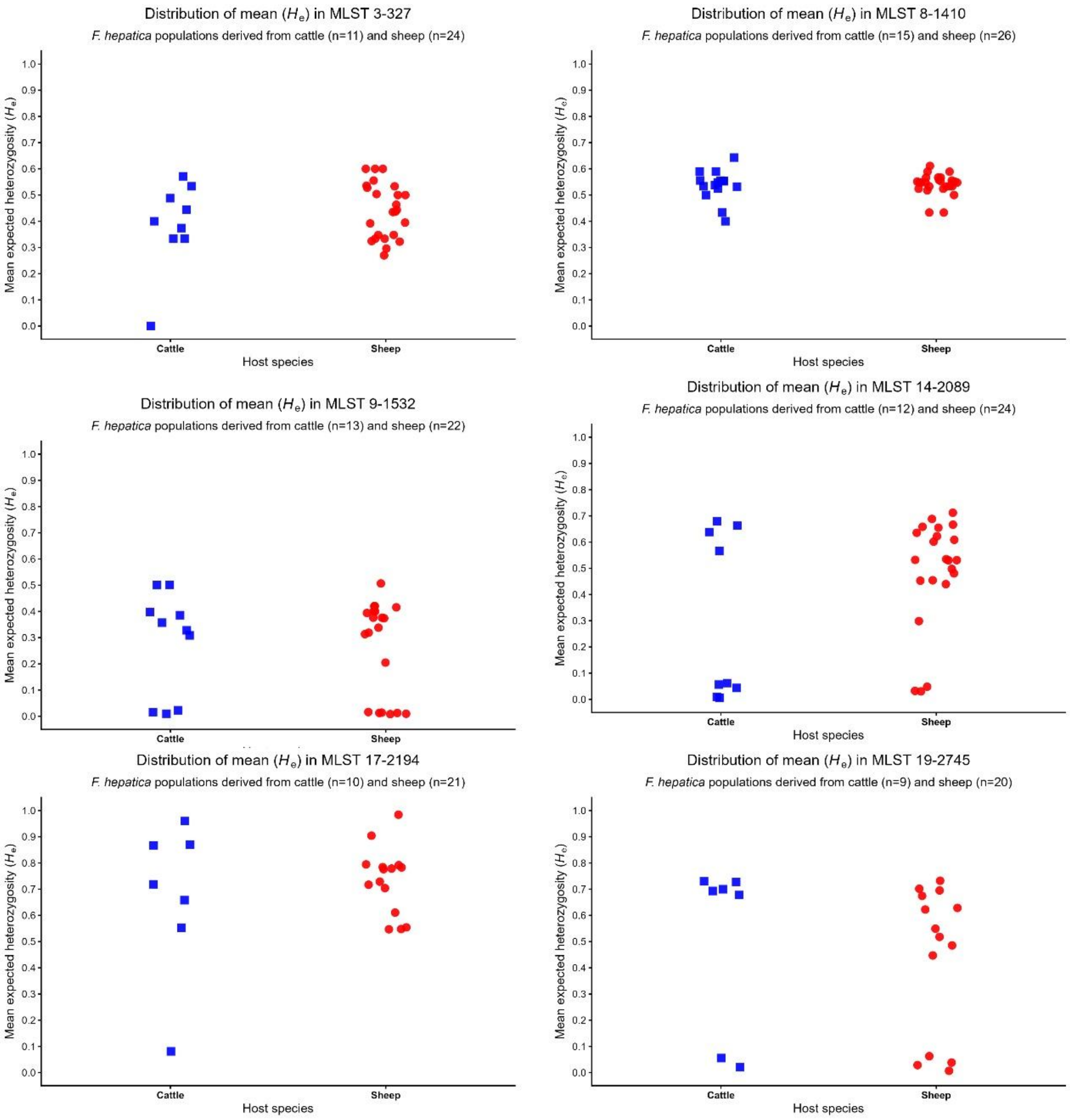


**Supplementary Fig. 3.** Distribution of mean expected heterozygosity (He) in cattle- and sheep-derived *F. hepatica* populations based on six MLST loci. Each square is represents cattle and dot shows sheep mean He value of an individual population. The y-axis indicates mean He values for cattle and sheep-derived populations, while the x-axis represents the host species.
