## Supplementary Table 1 for "Development and validation of a multilocus sequence typing scheme for *Fasciola hepatica* using next-generation deep amplicon sequencing"

**Supplementary Table 1:** *F. hepatica* populations and samples details. Samples collected within similar time frames from the same sheep or cattle farms, and veterinary practitioners were classified as a single parasite population.

| Population name | Samples included | Dates of sample collection | Sample type | County | Region | Host animal |
|---|---|---|---|---|---|---|
| P1C | SNo14<br>SNo15<br>Sno12 | 28/02/2023 | Faecal | Wiltshire | South West England | Cattle |
| P2C | SNO38 | 28/03/2023 | Faecal | Cheshire | North West England | Cattle |
| P3C | Sno44<br>Sno47 | 28/02/2023 | Faecal | Cumbria | North West England | Cattle |
| P4C | SNo383 | 26/03/2024 | Faecal | Cumbria | North West England | Cattle |
| P5S | Sno24 | 01/03/2023 | Faecal | Cumbria | North West England | Sheep |
| P6S | Sno376 | 22/03/2024 | Faecal | Cumbria | North West England | Sheep |
| P7S | SNo75 | 31/03/2023 | Faecal | Cumbria | North West England | Sheep |
| P8S | SNo74 | 28/03/2023 | Faecal | Staffordshire | West Midlands England | Sheep |
| P9S | SNo230<br>SNo231<br>SNo232<br>SNo267<br>SNo269 | 19/11/2023<br>19/11/2023<br>19/11/2023<br>13/12/2023<br>13/12/2023 | Faecal | Scottish Borders | Scottish Borders | Sheep |
| P10C | SNo241 | 28/11/2023 | Faecal | Peeblesshire | Scottish Borders | Cattle |
| P11S | SNo322<br>SNo323<br>SNo324<br>SNo325 | 02/02/2024 | Faecal | Scottish Borders | Scottish Borders | Sheep |
| P12S | SNo297<br>SNo298<br>SNO300 | 23/01/2024 | Faecal | Peeblesshire | Scottish Borders | Sheep |
| P13C | SNo326<br>SNo328<br>SNo330<br>SNo331 | 04/02/2024 | Faecal | Peeblesshire | Scottish Borders | Cattle |
| P14C | SNo407<br>SNo408 | 25/04/2024 | Faecal | Peeblesshire | Scottish Borders | Cattle |
| P15C | SNo283<br>SNo284<br>SNo285<br>SNo286<br>SNo287 | 04/01/2024 | Faecal | West Lothian | Southeastern Scotland | Cattle |

| | | | | | | |
|---|---|---|---|---|---|---|
| | SNo288 | | | | | |
| P16C | SNo301<br>SNo302<br>SNo303<br>SNo304<br>SNo305<br>SNo306 | 30/01/2024 | Faecal | Dorset | South West England | Cattle |
| P17C | SNo345<br>SNo346<br>SNo347<br>Sno349 | 15/02/2024 | Faecal | Dorset | South West England | Cattle |
| P18C | Sno341<br>Sno343<br>Sno344 | 15/02/2024 | Faecal | Dorset | South West England | Cattle |
| P19S | SNo313<br>SNo314<br>SNo316<br>SNo318 | 01/02/2024 | Faecal | South Lanarkshire | Southern Scotland | Sheep |
| P20S | SN0332 | 01/02/2024 | Faecal | South Lanarkshire | Southern Scotland | Sheep |
| P21S | SNo333 | 01/02/2024 | Faecal | South Lanarkshire | Southern Scotland | Sheep |
| P22S | SNo334<br>SNo335 | 01/02/2024 | Faecal | South Lanarkshire | Southern Scotland | Sheep |
| P23S | SNo336 | 01/02/2024 | Faecal | South Lanarkshire | Southern Scotland | Sheep |
| P24S | SNo338 | 01/02/2024 | Faecal | South Lanarkshire | Southern Scotland | Sheep |
| P25S | SNo289<br>SNo290<br>SNo291 | 01/10/2024 | Faecal | South Lanarkshire | Southern Scotland | Sheep |
| P26S | SNo390<br>SNo391<br>SNo392<br>SNo393<br>SNo394 | 18/04/2024 | Faecal | Essex | East of England | Sheep |
| P27S | Sno33<br>Sno36 | 25/03/2023 | Faecal | Devon | South West England | Sheep |
| P28C | Sno371 | 11/03/2024 | Faecal | Devon | South West England | Cattle |
| P29C | Sno5 | 14/02/2023 | Faecal | Gloucestershire | South West England | Cattle |
| P30S | MR-1 | 20/03/2023 | Worms | Tyrone Northern Ireland | Northern Ireland | Sheep |
| P31C | VPC-1 | 05/05/2023 | Worms | West Sussex | South East England | Cattle |
| P32S | aki1_sheep | 05/07/2023 | Worms | West Sussex | South East England | Sheep |
| P33C | Downlands-1 | 21/03/2024 | Worms | East Sussex | South East England | Cattle |

| | | | | | | |
|---|---|---|---|---|---|---|
| P34S | VPC-2 | 04/01/2024 | Worms | Kent | South East England | Sheep |
| P35S | FZY160i | 23/10/24 | Worms | Derbyshire | East Midlands England | Sheep |
| P36S | FZY160ii | 23/10/24 | Worms | Derbyshire | East Midlands England | Sheep |
| P37S | aki2 | 13/09/2024 | Worms | Renfrewshire, Scotland | West of Scotland | Sheep |
| P38S | aki3 | 20/09/2024 | Worms | Renfrewshire, Scotland | West of Scotland | Sheep |
| P39S | aki4 | 20/09/2024 | Worms | Renfrewshire, Scotland | West of Scotland | Sheep |
| P40S | aki5 | 15/11/2024 | Worms | Renfrewshire, Scotland | West of Scotland | Sheep |
| P41S | aki6 | 15/11/2024 | Worms | Renfrewshire, Scotland | West of Scotland | Sheep |
| P42S | SNo22 | 03/01/2023 | Faecal | Cumbria | North West England | Sheep |

Note: Veterinary practitioners' and farmers' names are not mentioned. However, Sno60 and SNo32 did not generate sequence reads after filtration, so these two samples were not included in the populations.
