## Supplementary Table 2 for "Development and validation of a multilocus sequence typing scheme for *Fasciola hepatica* using next-generation deep amplicon sequencing"

**Supplementary Table 2:** Primers designed to amplify regions in the *F. hepatica* reference genome (Fasciola_10x_pilon, GCA_900302435.1) containing 5 or more consecutive or non-consecutive SNPs.

| Primer ID showing Primer no./ Scaffold no. | Primer sequence | Product length (bp) | Estimated Tm | Scaffold ID and nucleotide number of extracted fragments from Ref. Genome based on 5 or more SNPs |
|---|---|---|---|---|
| 1/118 | (1 F) 5'-GAGCCTCAGCGTGAATTAGT-3'<br>(193 R) 5'-AGCCTTGACCCCAAAATTCC-3' | 193 | 57.7<br>58.4 | LT997055.1:246118-246322 |
| 2/302 | (7 F) 5'-CCGGTAAAAAGACGTCACCAC-3'<br>(194 R) 5'-CTTCGAAGTGGTATGAAGCTGT-3' | 189 | 59.5<br>58.4 | LT997239.1:1554389-1554593 |
| 3/327 | (18 F) 5'-ACCATATCACCAAGTGGAGCG-3'<br>(193 R) 5'-TCCCCAAATCTTCGGAGGTTC-3' | 176 | 60.1<br>59.7 | LT997264.1:51799-52003 |
| 4/331 | (11F) 5'-CGGCAATTGGATGGTGCAAA-3'<br>(187 R) 5'-GTCTGGCCTTGCGTCAAATG-3' | 177 | 60<br>60.1 | LT997268.1:711505-711709 |
| 5/388 | (4 F) 5'-TGTCGGACTATCTGAGAAAAGGA-3'<br>(193 R) 5'-ACACATGCGCACCGATAGAT-3' | 180 | 58.7<br>59.9 | LT997325.1:277029-277233 |
| 6/673 | (17F) 5'-ATCTCGTCCGATCACCACTG-3'<br>(192 R) 5'-GACTTTTGTCGTGTAGCCATAAA-3' | 176 | 59.3<br>57.4 | LT997610.1:704815-705019 |
| 7/1221 | (16 F) 5'-AGTTTCTGACCTTTTTGTCGTGT-3'<br>(193 R) 5'-TCAGCCAAAAACCACACGAAG-3' | 178 | 59<br>59.6 | LT998158.1:2994898-2995102 |
| 8/1410 | (3 F) 5'-GTGCCCAACCACGTTTTCAG-3'<br>(195 R) 5'-ATCAAACGGCGAAGTGAGGA-3' | 193 | 60.2<br>59.7 | LT998347.1:319687-319891 |
| 9/1532 | (11 F) 5'-GTTTGCCACAGGACTGGGAA-3'<br>(186 R) 5'-GAGCTGGGCAAAGCATAGGA-3' | 176 | 60.5<br>60.1 | LT998469.1:3904782-3904986 |
| 10/1632 | (27 F) 5'-ACGCACAATGAAAGAAGACACG-3'<br>(202 R) 5'-ACCTAGTAAGTGTTCAGTACCGT -3' | 176 | 60<br>59.5 | LT998569.1:251558-251762 |
| 11/1914 | (10 F) 5'-CCTCATCGACTAGTGGTGCC-3'<br>(194 R) 5'-TCAGTTCACTTTTGATCCGTGT-3' | 185 | 59.9<br>58.2 | LT998851.1:248913-249117 |
| 12/2022 | (23 F) 5'-CCCGATTCCCTCAACCTGTT-3' | 181 | 59.7 | LT998959.1:636703-636907 |

| | | | | |
|---|---|---|---|---|
| | (203 R) 5'-GTTCCCATGCCTCCACGTAC-3' | | 60.7 | |
| 13/2086 | (6 F) 5'-TGCCAAAGTTTGCTCAACGTT-3'<br>(201 R) 5'-CGAGTCAGGGAAGCTCAACA-3' | 196 | 59.8<br>59.7 | LT999023.1:846951-847155 |
| 14/2089 | (8 F) 5'-GACCAGTCCACTTTGACAAACTA-3'<br>(192 R) 5'-CGTCTCGCATGTACGACGTA-3' | 185 | 58.6<br>60 | LT999026.1:3794694-3794898 |
| 15/2089 | (22 F) 5'-TGTCTCTCGGATGTTGATCGT-3'<br>(199 R) 5'-ACTTCAAGTATCACGAGAGCTGA-3' | 178 | 58.9<br>59.2 | LT999026.1:5646988-5647192 |
| 16/2093 | (30 F) 5'-ATCACCGTCCGTGTTTTTCC-3'<br>(204 R) 5'-CCAATTACGTGTGATTCGTTTCG-3' | 175 | 58.8<br>58.9 | LT999030.1:1478846-1479050 |
| 17/2194 | (5 F) 5'-TAGCCCCTCACGTGGGTAAT-3'<br>(179 R) 5'-ACGTTATTATCATAGGCGATAGATGGT-3' | 175 | 60.3<br>59.9 | LT999131.1:399804-400008 |
| 18/2351 | (21 F) 5'-AGCTGAATAAGAGGCAAGAGGT-3'<br>(204 R) 5'-CAGACAGGCTAATGGAAAGTAGT-3' | 184 | 59.2<br>57.9 | LT999288.1:1153480-1153684 |
| 19/2745 | (14 F) 5'-GCCTCTGACTCAAATCGATCCT-3'<br>(191 R) 5'-ACTACTTCCCTTCGACCACCT-3' | 178 | 59.9<br>60.2 | LT999682.1:306828-307032 |
| 20/2815 | (1 F) 5'-CGCGGTAAAGGACAGTGATAAC-3'<br>(190 R) 5'-CTGCATAGTGGGGTACGTCG-3' | 190 | 59.4<br>60.2 | LT999752.1:2940358-2940562 |
