## Supplementary Table 5 for "Development and validation of a multilocus sequence typing scheme for *Fasciola hepatica* using next-generation deep amplicon sequencing"

**Supplementary Table. 5.** Sample details and quality assessment of DNA isolated from adult worms and selected for whole genome sequencing

| **Method** | **VPC-1 (ng/µl)** | **VPC-1 worm 2/3 (ng/µl)** | **VPC-1 worm 3/3 (ng/µl)** | **MR-1 (ng/µl)** | **MR-1 3/3 (in 70% ethanol) (ng/µl)** |
|---|---|---|---|---|---|
| Biodrop reader | **291.6** | 12.92 | 10.92 | **194.1** | 135 |
| $OD_{260/280}$ values | **1.928** | 1.891 | 1.752 | **1.832** | 1.821 |
| Qubit reader average | **135.0** | 2.04 | 0.925 | **66.80** | 30.5 |
| Host animal | Sheep (Steyning, West Sussex) | | | Cattle (Co Tyrone, Northern Ireland) | |

Note The samples highlighted in bold were selected for whole genome sequencing
