## Supplementary Table 6 for "Development and validation of a multilocus sequence typing scheme for *Fasciola hepatica* using next-generation deep amplicon sequencing"

**Supplementary Table 6:** Mapping data after alignment with reference sequences of *F. hepatica* and *F. gigantica*

| Sample ID | Host animal of sample | Q 30 score after rFASTp | Q 20 score after rFASTp | Ref. Seq. accession number | Ref. seq. owner name | Host animal of ref. seq. | N50 for ref. seq. | Parasite species | Overall alignment rate* | Mapping Percentage ** |
|---|---|---|---|---|---|---|---|---|---|---|
| VPC-1 | Cattle | 81.07% | 94.31% | GCA_002763495.2 | The Genome Institute USA (2019) | Sheep | 161.1 kb | *F. hepatica* | 88.89% | 99.90% |
| | | | | GCA_900302435.1 | University of Liverpool UK (2021) | Sheep | 1.9 Mb | *F. hepatica* | 93.35% | 99.99% |
| | | | | GCA_948099385.1_FhHiC23_genomic.fna *** | University of Liverpool UK (2023) | Sheep | 9.8 Mb | *F. hepatica* | 95.39% | 100% |
| MR-1 | Sheep | 90.74% | 99.67% | GCA_002763495.2 | The Genome Institute USA (2019) | Sheep | 161.1 kb | *F. hepatica* | 90.20% | 99.91% |
| | | | | GCA_900302435.1 | University of Liverpool UK (2021) | Sheep | 1.9 Mb | *F. hepatica* | 94.90% | 99.99% |
| | | | | GCA_948099385.1_FhHiC23_genomic.fna | University of Liverpool UK (2023) | Sheep | 9.8 Mb | *F. hepatica* | 96.57% | 100% |

*Overall alignment was determined by Bowtie2 **Mapping percentages were accessed in samtools using the HPC cluster at University of Surrey. ***GCA_948099385.1_FhHiC23 has now been replaced by improved version GCA_948099385.2
