## Supplementary Table 7 for "Development and validation of a multilocus sequence typing scheme for *Fasciola hepatica* using next-generation deep amplicon sequencing"

**Supplementary Table 7:** Summary of primer performance on DNA isolated from *F. hepatica* eggs

| **Primer ID/Scaffold No** | **Sample no 75 (Cumbria, Sheep)** EPG=(Cd/Fh (1/120)) | **Sample no 285 (Scottish Borders, Sheep)** EPG=Fh (15) | **Sample No 298 (Peeblesshire, Sheep)** EPG=Fh (4) | **Sample 316 (South Lanarskshire, Sheep)** EPG= Fh (18) | **Sample No 345 (West Dorset, Cattle)** EPG=Fh (96) |
|---|---|---|---|---|---|
| 1/118 | Band observed | -- | -- | -- | Band observed |
| 2/302 | Band observed (multiple) | Band observed | -- | -- | -- |
| 3/327 | Band observed (multiple) | Band observed | Band observed | Band observed | Band observed |
| 4/331 | -- | -- | -- | -- | -- |
| 5/388 | Band observed (multiple) | -- | -- | -- | -- |
| 6/673 | Band observed (multiple) | Band observed | Band observed | Band observed | Band observed |
| 8/1410 | Band observed | Band observed | Band observed | Band observed | Band observed |
| 9/1532 | Band observed | Band observed | Band observed | Band observed | Band observed |
| 10/1632 | Band observed | -- | Band observed | Band observed | Band observed |
| 11/1914 | Band observed (multiple) | Band observed (multiple) | Band observed (multiple) | Band observed | Band observed |
| 12/2022 | Band observed | -- | Band observed (multiple) | Band observed | Band observed |
| 13/2086 | -- | -- | -- | -- | -- |
| 14/2089 | Band observed | Band observed | Band observed (multiple) | Band observed | Band observed |
| 15/2089 | Band observed (multiple) | -- | Band observed (multiple) | Band observed | Band observed |
| 16/2093 | -- | -- | -- | Band observed (multiple) | Band observed |
| 17/2194 | -- | | Band observed | Band observed | Band observed |

| | | | | | |
|---|---|---|---|---|---|
| 19/2745 | Band observed (multiple) | Band observed | Band observed (multiple) | Band observed | Band observed (multiple) |
| 20/2815 | Band observed | Band observed | Band observed (multiple) | Band observed | -- |
| **Total** | **multiple 14** | **multiple9** | **multiple 12** | **multiple13** | **multiple 13** |
